# Arterial smooth muscle cell PKD2 (TRPP1) channels control systemic blood pressure: Arterial myocyte PKD2 controls blood pressure

**DOI:** 10.1101/320200

**Authors:** Simon Bulley, Carlos Fernandez-Pena, Raquibul Hasan, M. Dennis Leo, Padmapriya Muralidharan, Charles E. Mackay, Kirk W. Evanson, Sarah K. Burris, Qian Wang, Korah P. Kuruvilla, Jonathan H. Jaggar

## Abstract

Systemic blood pressure is determined, in part, by arterial smooth muscle cells (myocytes). Several Transient Receptor Potential (TRP) channels are proposed to be expressed in arterial myocytes, but it is unclear if these proteins control physiological blood pressure and contribute to hypertension *in vivo*. We generated the first inducible, smooth muscle-specific knockout for a TRP channel, namely for PKD2 (TRPP1), to investigate arterial myocyte and blood pressure regulation by this protein. Using this model, we show that intravascular pressure and α_1_-receptors activate PKD2 channels in arterial myocytes of different systemic organs. PKD2 channel activation in arterial myocytes leads to an inward Na+ current, membrane depolarization and vasoconstriction. Inducible, smooth muscle cell-specific PKD2 knockout lowers both physiological blood pressure and hypertension and prevents pathological arterial remodeling during hypertension. In summary, we show for the first time that arterial myocyte PKD2 channels control systemic blood pressure and targeting reduces high blood pressure.

## Introduction

Systemic blood pressure is controlled by total peripheral resistance, which is determined by the diameter of small arteries and arterioles. Arterial smooth muscle cell (myocyte) contraction reduces luminal diameter, leading to an increase in systemic blood pressure, whereas relaxation results in vasodilation that decreases blood pressure. Membrane potential is a primary determinant of arterial contractility^1^. Depolarization activates voltage-dependent calcium (Ca^2+^) channels (Ca_v_) in myocytes, leading to an increase in intracellular Ca^2+^ concentration [Ca^2+^]_i_ and vasoconstriction. In contrast, hyperpolarization reduces [Ca^2+^]i, resulting in vasodilation^1^. *In vitro* studies have identified several different types of ion channel that regulate arterial myocyte contractility. However, the molecular identity of arterial myocyte ion channels that regulate blood pressure and their mechanisms of modulation *in vivo* poorly understood. Given that hypertension is associated with altered arterial contractility, myocyte ion channels that contribute to high blood pressure are also important to investigate.

Transient Receptor Potential (TRP) channels constitute a family of ~28 proteins that are subdivided into 6 different classes, including polycystin (TRPP), canonical (TRPC), vanilloid (TRPV), ankyrin (TRPA), and melastatin (TRPM) ^2^. Studies performed using cultured and non-cultured cells and whole arteries, which contain many different cell types, have suggested that approximately thirteen different TRP channels may be expressed in arterial myocytes, including PKD2 (also termed TRPP1), TRPC1, TRPC3-6, TRPV1-4, TRPA1, TRPM4 and TRPM8 ^2^. In many cases TRP channel expression was reported in vasculature that does not control systemic blood pressure, including conduit vessels, cerebral arteries, portal vein and pulmonary arteries ^2^. It is unclear whether the same TRP channels are expressed in arterial myocytes of different organs, if TRP proteins present are similarly regulated or if they perform analogous physiological functions. Thus, the extrapolation of findings in one arterial preparation to other blood vessels appears inappropriate, requiring targeted approaches, such as cell-specific genetic knockouts. Blood pressure measurements in constitutive, global TRPC6, TRPM4 and TRPV4 channel knockout mice produced inconsistent results or generated complex findings that were associated with compensatory mechanisms ^3–6^. Thus which, if any, arterial myocyte TRP channels control blood pressure is unclear. In this study, we generated the first inducible, myocyte-specific knockout mouse for any TRP channel, specifically for PKD2, to investigate blood pressure regulation by this protein.

PKD2 is a six transmembrane domain protein with cytoplasmic N and C termini ^7^. PKD2 is expressed in myocytes of rat and human cerebral arteries, mouse and human mesenteric arteries and in porcine whole aorta ^8–10^. RNAi-mediated knockdown of PKD2 inhibited pressure-induced vasoconstriction (myogenic tone) in cerebral arteries ^10 11^. In global, constitutive PKD2+/− mice, an increase in actin and myosin expression lead to larger phenylephrine-induced contractions in aorta and mesenteric arteries ^12, 13^. It has also been proposed that arterial myocyte PKD2 channels inhibit myogenic tone in mesenteric arteries ^11^. Thus, *in vitro* studies have generated variable findings regarding physiological functions of arterial myocyte PKD2 channels.

Here, our data show that inducible and cell-specific PKD2 channel knockout in systemic artery myocytes reduces systemic blood pressure. We demonstrate that intravascular pressure and α1-adrenoceptor activation activate arterial myocyte PKD2 channels in systemic arteries of different tissues, leading to an inward sodium current (I_Na_), membrane depolarization and vasoconstriction. We also show that PKD2 channels are upregulated during hypertension and that PKD2 knockout in myocytes causes vasodilation, attenuates remodeling of the arterial wall and reduces high blood pressure during hypertension. In summary, our study indicates that PKD2 channels in systemic artery myocytes control physiological blood pressure and are upregulated during hypertension, contributing to the blood pressure elevation.

## Methods

**Animals.** All procedures were approved by the Animal Care and Use Committee of the University of Tennessee. PKD2^fl/fl^ mice with loxP sites flanking exons 11–13 of the *Pkd2* gene were obtained from the John Hopkins PKD Core. TRPP1^fl/fl^ mice were crossed with tamoxifen-inducible smooth muscle-specific Cre mice (SMMHC-CreER^T2^, Jackson Labs, ref. ^36^) to produce PKD2^fl/fl^:smCre^+^ mice. Male mice (6–10 weeks of age) were injected with tamoxifen (1 mg/ml, i.p.) once per day for 5 days and studied no earlier than 17 days after the last injection. C57BL/6J mice (12 weeks old) were purchased from Jackson Laboratories. Angiotensin II (1.5 ng/g/min) and saline (0.9 NaCl) were infused in mice using subcutaneous osmotic minipumps (Alzet).

**Tissue preparation and myocyte isolation.** Mice were euthanized with isofluorane (1.5 %) followed by decapitation. First- to fifth-order mesenteric and hindlimb (saphenous, popilital and gastrocnemius) arteries were removed and placed into ice-cold physiological saline solution (PSS) that contained (in mmol/L): 112 NaCl, 6 KCl, 24 NaHCO_3_, 1.8 CaCl_2_, 1.2 MgSO_4_, 1.2 KH_2_PO_4_ and 10 glucose, gassed with 21% O_2_, 5% CO_2_ and 74% N_2_ to pH 7.4. Arteries were cleaned of adventitial tissue and myocytes dissociated in isolation solution containing (in mmol/L): 55 NaCl, 80 sodium glutamate, 5.6 KCl, 2 MgCl_2_, 10 HEPES and 10 glucose (pH 7.4, NaOH) using enzymes, as previously described ^39^.

**Genomic PCR.** Genomic DNA was isolated from mesenteric and hindlimb arteries using a Purelink Genomic DNA kit (ThermoFisher Scientific). Reaction conditions used are outlined in the Baltimore PKD core center genotyping protocol (http://baltimorepkdcenter.org/mouse/PCR%20Protocol%20for%20Genotyping%20PKD2KO%20and%20PKD2%5Eneo.pdf). Briefly, genotyping was performed using a 3-primer strategy, with primers a (5’- CCTTTCCTCTGGTTCTGGGGAG), b (5’ -GTTGATGCTTAGCAGATGATGGC) and c (5’- CTGACAGGCACCTACAGAACAGTG) used to identify floxed and deleted alleles (Supplemental Figure 1).

**RT-PCR.** Fresh, dissociated mesenteric artery myocytes were manually collected using an enlarged patch pipette under a microscope. Total RNA was extracted from ~500 myocytes using the Absolutely RNA Nanoprep kit (Agilent Technologies, Santa Clara, CA, USA). First-strand cDNA was synthesized from 1–5 ng RNA using SuperScript IV (Invitrogen, Life Technologies). PCR was performed on first-strand cDNA using the following conditions: an initial denaturation at 94°C for 2 min, followed by 40 cycles of denaturation at 94°C for 30 s, annealing at 56°C for 30 s, and extension at 72°C for 1 min. PCR products were separated on 2 % agarose–TEA gels. Primers were used to amplify transcripts of PKD2, aquaporin 4, myosin heavy chain 11, platelet-endothelial cell adhesion molecule 1 (PECAM-1) and actin (Supplemental Table 1). The PKD2 forward primer spanned the junction of exons 9 and 10 and the reverse primer annealed to exon 13.

**Western blotting.** Proteins were separated on 7.5% SDS-polyacrylamide gels and blotted onto PVDF membranes. Membranes were blocked with 5% milk and incubated with the following primary antibodies: Ca_v_1.2 (Neuromab), PKD1 and PKD2 (Santa Cruz), ANO1, TRPC6 and TRPM4 (Abcam) and actin (MilliporeSigma) overnight at 4°C. Membranes were washed and incubated with horseradish peroxidase-conjugated secondary antibodies at room temperature. Protein bands were imaged using an Amersham Imager 600 gel imaging system (GE Healthcare) and quantified using ImageJ software.

**En-face arterial immunofluorescence.** Arteries were cut longitudinally and fixed with 4 % paraformaldehyde in PBS for 1 h. Following a wash in PBS, arteries were permeabilized with 0.2 % Triton X-100, blocked with 5 % goat serum and incubated overnight with PKD2 primary antibody (Baltimore PKD Center) at 4 °C. Arteries were then incubated with Alexa Fluor 555 rabbit anti-mouse secondary antibody (1:400; Molecular Probes) and 4’,6-diamidino-2-phenylindole, dihydrochloride (DAPI) (1:1000; Thermo Scientific) for 1 hr at room temperature. Segments were washed with PBS, oriented on slides with the endothelial layer downwards and mounted in 80 % glycerol solution. DAPI and Alexa 555 were excited at 350 nm and 555 nm with emission collected at ≤437 nm and ≥555 nm, respectively.

**Isolated arterial myocyte immunofluorescence.** Myocytes were plated onto poly-L-lysine-coated coverslips, fixed with 3.7 % paraformaldehyde in PBS and permeabilized with 0.1 % Triton X-100. After blocking with 5 % BSA, cells were incubated with mouse monoclonal anti-PKD2 antibody (Santa Cruz) overnight at 4 °C. Slides were then washed and incubated with Alexa Fluor 555 rabbit anti-mouse secondary antibody (Molecular Probes). Secondary antibodies were washed and coverslips mounted onto slides. Images were acquired using a Zeiss 710 (Carl Zeiss) laser-scanning confocal microscope and 40x and 63x oil immersion objectives.

**Kidney histology.** Kidney sections were stained with H&E and examined by Probetex, Inc (San Antonio, Texas). Briefly, tubules, glomeruli and vasculature were examined for frequency or homogeneity of pathologic abnormalities. These included characteristics of hypercellularity, hypocellularity, necrosis, apoptosis, matrix accumulation, inflammation, fibrosis and protein droplets. The size and thickness of the cortex, medulla, papilla, glomeruli, tubules and vasculature were also examined.

**Telemetric blood pressure measurements.** Radiotelemetric transmitters (PA-C10, Data Sciences International) were implanted subcutaneously into anesthetized mice, with the sensing electrode placed in the aorta via the left carotid artery. Seven days later blood pressures were measured using a PhysioTel Digital telemetry platform (Data Sciences International). Dataquest A.R.T. software was used to acquire and analyze data.

**Echocardiography.** Mice were anesthetized with isofluorane. Ultrasound gel was placed on a hairless area of the chest before and echocardiography performed using a Visual Sonics Vevo 2100 system. Anesthetic depth, heart rate and body temperature were monitored throughout the procedure.

**Arterial histology.** Arteries were fixed with paraformaldehyde and embedded in paraffin. 5 μm thick sections were cut using a microtome and mounted on slides. Sections were de-paraffinized, blocked in BSA and incubated with H&E. Images were acquired using a transmitted light microscope (Nikon Optiphot-2) and measurements made using Stereo Investigator software (MicrobrightField, Inc.). Wall-to-lumen ratios were calculated as wall thickness/lumen diameter, where the wall (tunica media) thickness and lumen diameter of each section was the averages of four and two equidistant measurements, respectively.

**Blood and urine analysis.** Retro-orbital blood was drawn from isofluorane-anesthetized mice using a Microvette Capilliary Blood Collection System (Kent Scientific Corporation). Plasma was extracted and angiotensin II (Elabscience), aldosterone (Mybiosource) and atrial natriuretic peptide (Elabscience) concentrations measured using commercially available ELISA kits and an EL800 plate reader (BioTeK). Mice were housed in individual metabolic cages for 72 hours and urine collected for the final 24 hours. Plasma and urine electrolyte concentrations were measured using the MMPC Core at Yale University.

**Pressurized artery myography.** Experiments were performed using isolated mouse third-, fourth and fifth-order mesenteric arteries and first-order gastrocnemius arteries using PSS gassed with 21% O_2_/5% CO_2_/74% N_2_ (pH 7.4). Arterial segments 1–2 mm in length were cannulated at each end in a perfusion chamber (Living Systems Instrumentation) continuously perfused with PSS and maintained at 37°C. Intravascular pressure was altered using an attached reservoir and monitored using a pressure transducer. Luminal flow was absent during experiments. Arterial diameter was measured at 1 Hz using a CCD camera attached to a Nikon TS100-F microscope and the automatic edge-detection function of IonWizard software (Ionoptix). Myogenic tone was calculated as: 100 × (1-D_active_/D_passive_) where D_active_ is active arterial diameter, and D_passive_ is the diameter determined in the presence of Ca^2+^-free PSS supplemented with 5 mmol/L EGTA).

**Perfused hindlimb pressure measurements.** Isolated hindlimbs were inserted into a chamber containing gassed PSS (21% O_2_/5% CO_2_/74% N_2_) that was placed on a heating pad to maintain temperature at 37 °C. The femoral artery was cannulated with a similar diameter glass pipette and perfused with gassed PSS at 37°C using a peristaltic pump. Perfusion pressure was measured using a pressure transducer connected to the inflow. The flow rate was increased stepwise from 0 to 2.5 mL/min in 0.25 mL/min steps to generate a response curve. Values were corrected by subtracting the pressure produced by the pipette alone at each flow rate. Prior to measuring responses to phenylephrine, flow rate was adjusted to maintain a constant perfusion pressure of 80 mmHg. Data were recorded and analyzed using IonWizard software (Ionoptix).

**Wire myography.** Mesenteric artery segments (1st and 2nd order, 2 mm in length) were mounted on tungsten wires in a multi-channel myography system (Danish Myo Technology). Chambers contained PSS that was continuously gassed with 21% O_2_/5% CO_2_/74% N_2_ (pH 7.4). Arterial rings were subjected to a resting tension of 10 mN and allowed to equilibrate prior to experimentation. Responses were measured to increasing concentrations of phenylephrine or 60 mmol/L K^+^. Data were acquired and analyzed using LabChart software (ADInstruments).

**Pressurized artery membrane potential measurements.** Membrane potential was measured by inserting sharp glass microelectrodes (50–90 mΩ) filled with 3 mol/L KCl into the adventitial side of pressurized mesenteric or hindlimb arteries. Membrane potential was recorded using a WPI FD223a amplifier and digitized using a MiniDigi 1A USB interface, pClamp 9.2 software (Axon Instruments) and a personal computer. Criteria for successful intracellular impalements were: (1) a sharp negative deflection in potential on insertion; (2) stable voltage for at least 1 minute after entry; (3) a sharp positive voltage deflection on exit from the recorded cell and (4) a <10% change in tip resistance after the impalement.

**Patch-clamp electrophysiology.** Isolated arterial myocytes were allowed to adhere to a glass coverslip in a recording chamber. The conventional whole-cell configuration was used to measure non-selective cation currents (I_cat_) by applying either voltage ramps (0.13 mV/ms) or stepwise depolarizations between −100 mV and +100 mV from a holding potential of −40 mV. For cell swelling experiments, the pipette solution contained (in mmol/L): Na^+^ aspartate 115, mannitol 50, HEPES 10, glucose 10, EGTA 1, NaGTP 0.2, with free Mg^2+^ and Ca^2+^ of 1 mmol/L and 200 nmol/L, respectively (pH 7.2, NaOH). Isotonic (300 mOsm) bath solution contained (in mmol/L): Na^+^ aspartate 115, mannitol 50, glucose 10, HEPES 10, CaCl_2_ 2, MgCl_2_ 1 (pH 7.4, NaOH). Hypotonic (250 mOsm) bath solution was the same formulation as isotonic bath solution with the exclusion of mannitol (pH 7.4, NaOH). For experiments that measured I_Cat_ regulation by phenylephrine, the bath solution contained (in mmol/L): 140 NaCl, 10 glucose, 10 HEPES, 1 MgCl_2_, and pH was adjusted to 7.4 with NaOH. Pipette solution contained: 140 NaCl, 10 HEPES, 10 Glucose, 5 EGTA, 1 MgATP, 0.2 NaGTP, and pH was adjusted to 7.2 with NaOH. Total MgCl_2_ and total CaCl_2_ were adjusted to give free Mg^2+^ concentrations of 1 mmol/L and free Ca^2+^ of 200 nM. Free Mg^2+^ and Ca^2+^ were calculated using WebmaxC Standard (http://www.stanford.edu/~cpatton/webmaxcS.htm). Currents were recorded using an Axopatch 200B amplifier and Clampex 10.4 (Molecular Devices), digitized at 5 kHz and filtered at 1 kHz. Offline analysis was performed using Clampfit 10.4.

**Arterial biotinylation.** Procedures used were similar to those previously described ^40^. Briefly, arteries were biotinylated with EZ-Link Sulfo-NHS-LC-LC-Biotin and EZ-Link Maleimide-PEG2-Biotin. Unbound biotin was quenched with glycine-PBS, washed with PBS and then homogenized in radioimmunoprecipitation assay (RIPA) buffer. Protein concentration was normalized and biotinylated surface protein was captured by incubating cell lysates with avidin beads (Pierce) at 4°C. Proteins present in biotinylated and non-biotinylated samples were identified using Western blotting.

**Statistical analysis.** OriginLab and GraphPad InStat software were used for statistical analyses. Values are expressed as mean ± SEM. Student t-test was used for comparing paired and unpaired data from two populations and ANOVA with Student-Newman-Keuls post hoc test used for multiple group comparisons. P<0.05 was considered significant. Power analysis was performed to verify that the sample size gave a value of >0.8 if *P* was >0.05.

## Results

### Generation of tamoxifen-inducible smooth muscle-specific PKD2 knockout mice

Mice with loxP sites flanking exons 11 and 13 (PKD2^fl/fl^) of the *Pkd2* gene were crossed with tamoxifen-inducible SMMHC-CreERT2 mice, producing a PKD2^fl/fl^:smMHC-Cre^+^ line (Supplemental Figure 1A). Genotyping was performed using a 3-primer (a, b, c) strategy to identify wild-type (in C57BL/6J), floxed and deleted PKD2 alleles (Supplemental Figure 1B). PCR of genomic DNA from mesenteric and hindlimb arteries of wild-type mice that lack loxP sites produced a 232 bp transcript (Supplemental Figures 1B and 2). Tamoxifen-treated PKD2^fl/fl^ mouse arteries and aorta produced a transcript of 318 bp, which arose from primers a and b (Supplemental Figures 1B and 2). PCR amplified a 209 bp transcript in vasculature from tamoxifen-treated PKD2^fl/fl^:smCre^+^ mice due to primers a and c, confirming loss of the primer b annealing site. PCR of vasculature in PKD2^fl/fl^:smCre^+^ arteries also produced a faint 318 bp band, suggesting that PKD2 is expressed in vascular wall cell types other than myocytes where DNA would not undergo recombination (Supplemental Figure 2).

### PKD2 transcripts and protein are absent in arterial myocytes of tamoxifen-treated PKD2^fl/fl^:smCre^+^ mice

RT PCR was performed on RNA extracted from populations of myocytes (~500) that had been isolated using enzymes and individually harvested from mesenteric arteries of tamoxifen-treated PKD2^fl/fl^ and tamoxifen-treated PKD2^fl/fl^:smCre^+^ mice. Transcripts for both actin and myosin heavy chain 11, a smooth muscle-specific marker, were amplified by arterial myocyte cDNA from both genotypes (Figure 1A). In contrast, products for PECAM, an endothelial cell marker, and aquaporin 4, an astrocyte marker, were not amplified, suggesting that the isolated mRNA was from pure smooth muscle (Figure 1A). PKD2 primers were designed to anneal to a sequence in exons 9/10 (forward) and 13 (reverse), which span the recombination site (Supplemental Figure 1A). Transcripts for PKD2 were amplified by cDNA from arterial myocytes of PKD2^fl/fl^ mice, but not arterial myocytes of PKD2^fl/fl^:smCre^+^ mice (Figure 1A). These data indicate that tamoxifen induces PKD2 knockout in arterial myocytes of the PKD2^fl/fl^:smCre^+^ mice (Figure 1A).

**Figure 1:**
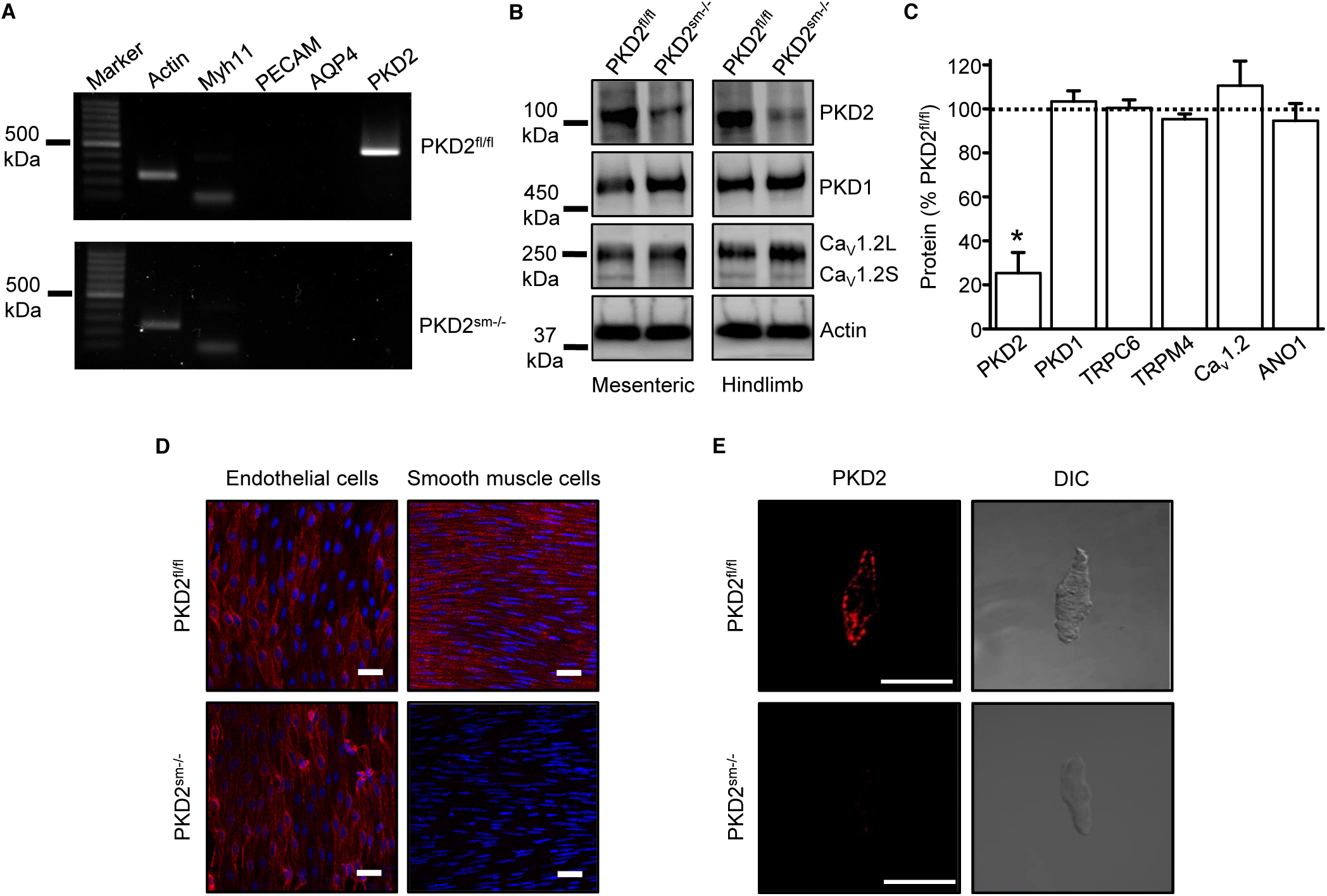
Activation of Cre recombinase abolishes PKD2 in arterial myocytes of PKD2^fl/fl^:smCre^+^ mice. **A.** RT-PCR showing the absence of a PKD2 transcript in isolated myocytes from tamoxifen-treated PKD2^fl/fl^:smCre^+^ mice**. B**: Western blots illustrating the effect of tamoxifen-treatment in PKD2^fl/fl^ and PKD2^fl/fl^smCre^+^ mice on PKD2, Ca_v_1.2L (full-length Ca_v_1.2) and Ca_v_1.2S (short Ca_v_1.2) proteins in mesenteric and hindlimb arteries. **C**: Mean data for proteins in mesenteric arteries of tamoxifen-treated PKD2^fl/fl^smCre^+^ mice when compared to those in tamoxifen-treated PKD2^fl/fl^ mice. n=4–5. * indicates p<0.05. **D**: *En-face* immunofluorescence imaging illustrating that PKD2 protein (red, Alexa Fluor 555) is abolished in myocytes of mesenteric arteries in tamoxifen-treated PKD2^fl/fl^:smCre^+^ mice (representative of 6 arteries). In contrast, PKD2 protein in endothelial cells is unaltered. Nuclear staining (DAPI) is also shown. Scale bars = 20 μm. **E**: Confocal and DIC images illustrating that PKD2 protein (Alexa Fluor 555) is abolished in isolated mesenteric artery myocytes of tamoxifen-treated PKD2^fl/fl^smCre^+^ mice (representative data from 5 PKD2^fl/fl^ and 5 PKD2^fl/fl^:smCre^+^ mice). Scale bars = 10 μm.

Western blotting was performed to quantify proteins in intact arteries of tamoxifen-injected PKD2^fl/fl^:smCre^+^ and tamoxifen-injected PKD2^fl/fl^ mice. In mesenteric and hindlimb arteries of PKD2^fl/fl^:smCre^+^ mice, PKD2 protein was ~25.3% and 32.6%, respectively of that in PKD2^fl/fl^ controls (Figures 1B, C, Supplemental Figures 3A and B). In contrast, TRPC6, TRPM4, and ANO1 channels and PKD1, which can form a complex with PKD2 ^14, 15^, were similar between genotypes (Figure 1C, Supplemental Figure 3A). Ca_v_1.2 protein was similar in mesenteric arteries, but slightly higher (~25.6 %) in hindlimb arteries of PKD2^sm−/−^ mice (Figures 1B, C, Supplemental Figure 3B). In aorta of PKD2^fl/fl^:smCre^+^ mice, PKD2 protein was ~46.8 % of that in PKD2^fl/fl^, whereas Ca_v_1.2 and PKD1 were similar (Supplemental Figures 3C, D). Immunofluorescence demonstrated that PKD2 protein was present in myocytes of intact arteries and isolated arterial myocytes of tamoxifen-treated PKD2^fl/fl^ control mice, but absent in myocytes of tamoxifen-treated PKD2^fl/fl^:smCre^+^ mice (Figures 1D, E). In contrast, PKD2 protein was present in endothelial cells of both tamoxifen-treated PKD2^fl/fl^ and PKD2^fl/fl^:smCre^+^ mouse arteries (Figure 1D). Thus, the remaining PKD2 protein detected in Western blots of tamoxifen-treated PKD2^fl/fl^:smCre^+^ mouse arteries is PKD2 present in cell types other than myocytes that would not be targeted by the smooth muscle-specific Cre. These data indicate that PKD2 is expressed in myocytes of mesenteric and hindlimb arteries and aorta and that tamoxifen treatment of PKD2^fl/fl^:smCre^+^ mice selectively abolishes PKD2 expression in myocytes. From this point in the manuscript, tamoxifen-treated PKD2^fl/fl^:smCre^+^ mice will be referred to as PKD2^sm−/−^. Tamoxifen-treated PKD2^fl/fl^ mice were used as controls in all experiments.

### PKD2^sm−/−^ mice are hypotensive

Telemetry indicated that diastolic and systolic blood pressures were both lower in PKD2^sm−/−^ mice than in PKD2^fl/fl^ mice (Figure 2A, 2B). Mean arterial pressure (MAP) was lower in PKD2^sm−/−^ mice during both day and night cycles, was sustained for days and on average was ~22.5 % lower in PKD2^sm−/−^ mice than in PKD2^fl/fl^ mice (Figure 2C, Supplemental Figure 4A). Locomotion was similar between genotypes, indicating that the lower blood pressure in PKD2^sm−/−^ mice was not due to inactivity (Supplemental Figure 4B). Echocardiography indicated that cardiac output, fractional shortening, ejection fraction and heart rate were similar in PKD2^sm−/−^ and PKD2^fl/fl^ mice (Figure 2D). PKD2^sm−/−^ kidney glomeruli and tubules were normal and indistinguishable from those of PKD2^fl/fl^ mice (Figure 2E). Plasma angiotensin II, aldosterone and ANP and plasma and urine electrolytes were similar in PKD2^sm−/−^ and PKD2^fl/fl^ mice (Table 1, p>0.05 for all). These data indicate that arterial myocyte PKD2 channels control systemic blood pressure and that cardiac function and kidney anatomy and function are similar in PKD2^sm−/−^ and PKD2^fl/fl^ mice.

**Figure 2:**
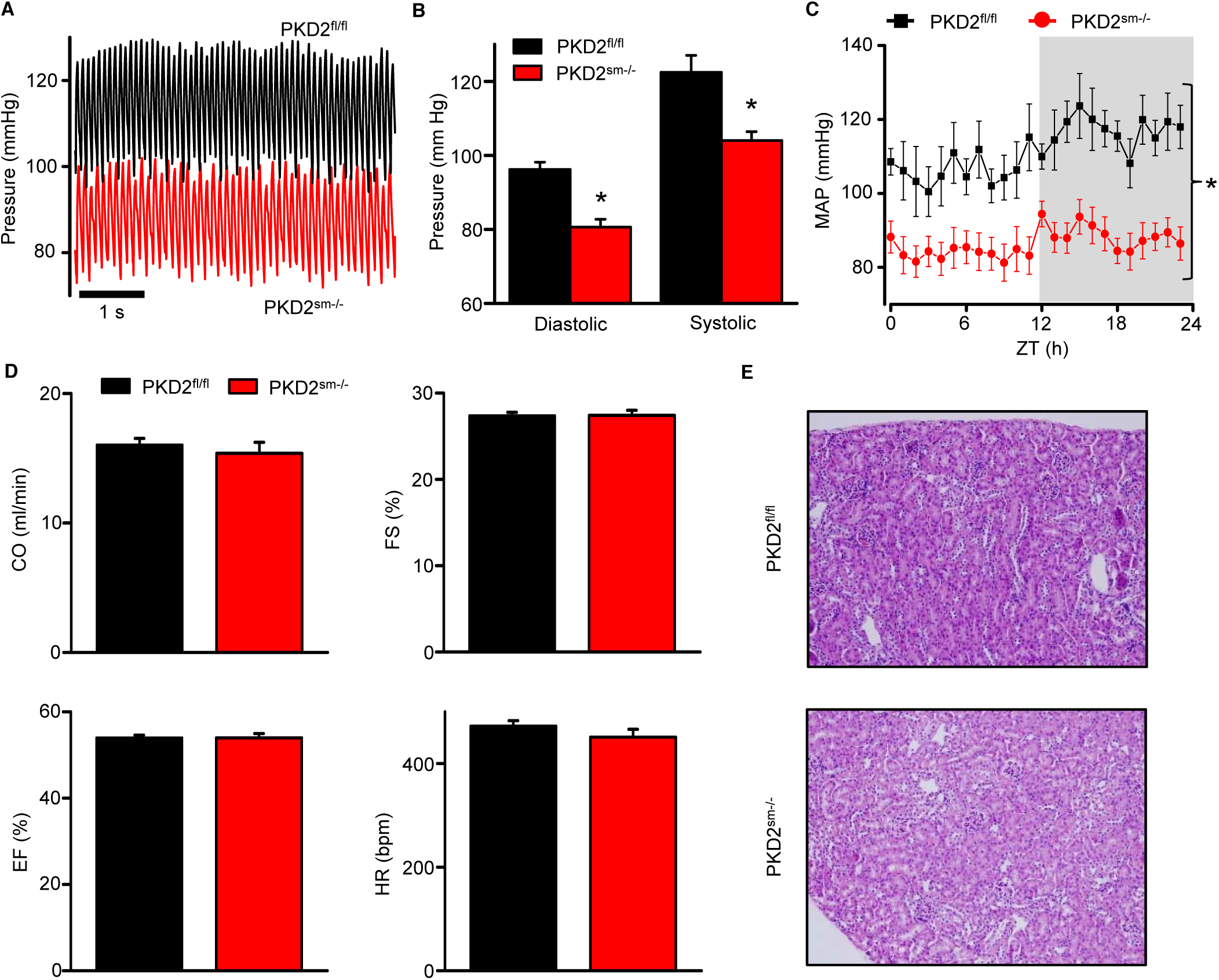
PKD2^sm−/−^ mice are hypotensive with normal cardiac function and renal histology. **A:** Original telemetric blood pressure recordings from PKD2^sm−/−^ and PKD2^fl/fl^ mice. **B**: Mean systolic and diastolic blood pressures in PKD2^fl/fl^ (n=11) and PKD2^sm−/−^ (n=12) mice. * indicates p<0.05 vs PKD2^fl/fl^. **C**: Mean arterial blood pressures (MAP) in PKD2^fl/fl^ (n=11) and PKD2^sm−/−^ (n=12) mice during day and night (gray) cycles. ZT: Zeitgeber Time. * indicates p<0.05 vs PKD2^fl/fl^ for all data points. **D**: Mean echocardiography data. Cardiac output (CO), fractional shortening (FS), ejection fraction (EF) and heart rate (HR). (PKD2^fl/fl^, n=5; PKD2^sm−/−^ mice, n=4). **E**: Representative images of H&E stained kidney cortex used for histological assessment (n=3 mice used for for each group).

**Table 1.**
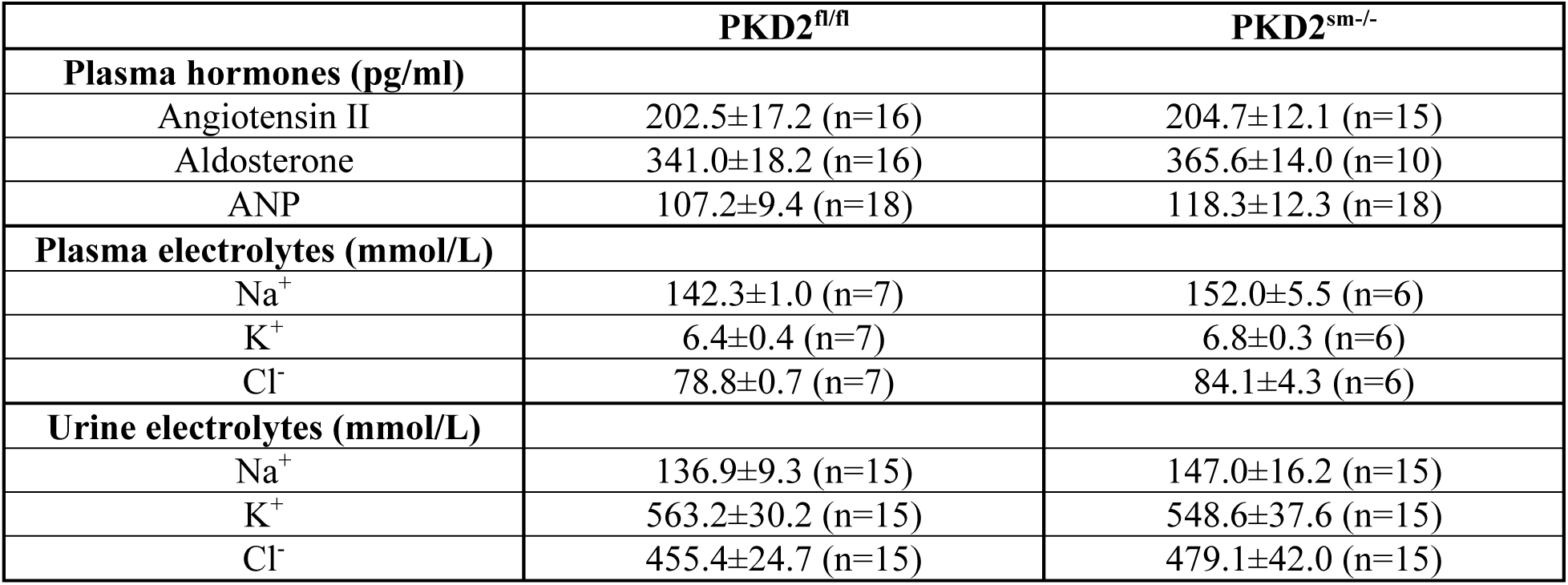
Plasma hormones and plasma and urine electrolytes.

### Myocyte PKD2 channels are essential for pressure-induced vasoconstriction in hindlimb arteries

Intravascular pressure stimulates vasoconstriction in small, resistance-size arteries. This myogenic response is a major regulator of both regional organ blood flow and systemic blood pressure. To determine whether PKD2 channels contribute to myogenic tone, pressure-induced (20–100 mmHg) vasoconstriction was measured in first-order gastrocnemius arteries. Intravascular pressures greater than 40 mmHg produced increasing levels of constriction in PKD2^fl/fl^ arteries, reaching ~20.9 % of passive diameter (tone) at 100 mmHg (Figures 3A, B). In contrast, pressure-induced vasoconstriction was robustly attenuated in PKD2^sm−/−^ gastrocnemius arteries, which developed only ~5.4 % tone at 100 mmHg or ~25.9 % of that in PKD2^fl/fl^ arteries (Figures 3A, B). The passive diameters of PKD2^fl/fl^ and PKD2^sm−/−^ gastrocnemius arteries were similar (Supplemental Figure 5A). Depolarization (60 mmol/L K^+^) stimulated slightly larger vasoconstriction in PKD2^sm−/−^ than PKD2^fl/fl^ gastrocnemius arteries (Supplemental Figure 5B). These data are consistent with the small increase in Ca_v_1.2 protein measured in gastrocnemius arteries of PKD2^sm−/−^ mice and indicate that: 1) attenuation of the myogenic response is not due to loss of voltage-dependent Ca^2+^ channel function and 2) that systemic blood pressure may control Ca_v_1.2 expression in hindlimb arteries, but not in mesenteric arteries or aorta (Supplemental Figure 3). In contrast to attenuated pressure-induced vasoconstriction, angiotensin II stimulated similar vasoconstrictions in gastrocnemius arteries of PKD2^sm−/−^ and PKD2^fl/fl^ mice (Supplemental Figure 5D).

**Figure 3:**
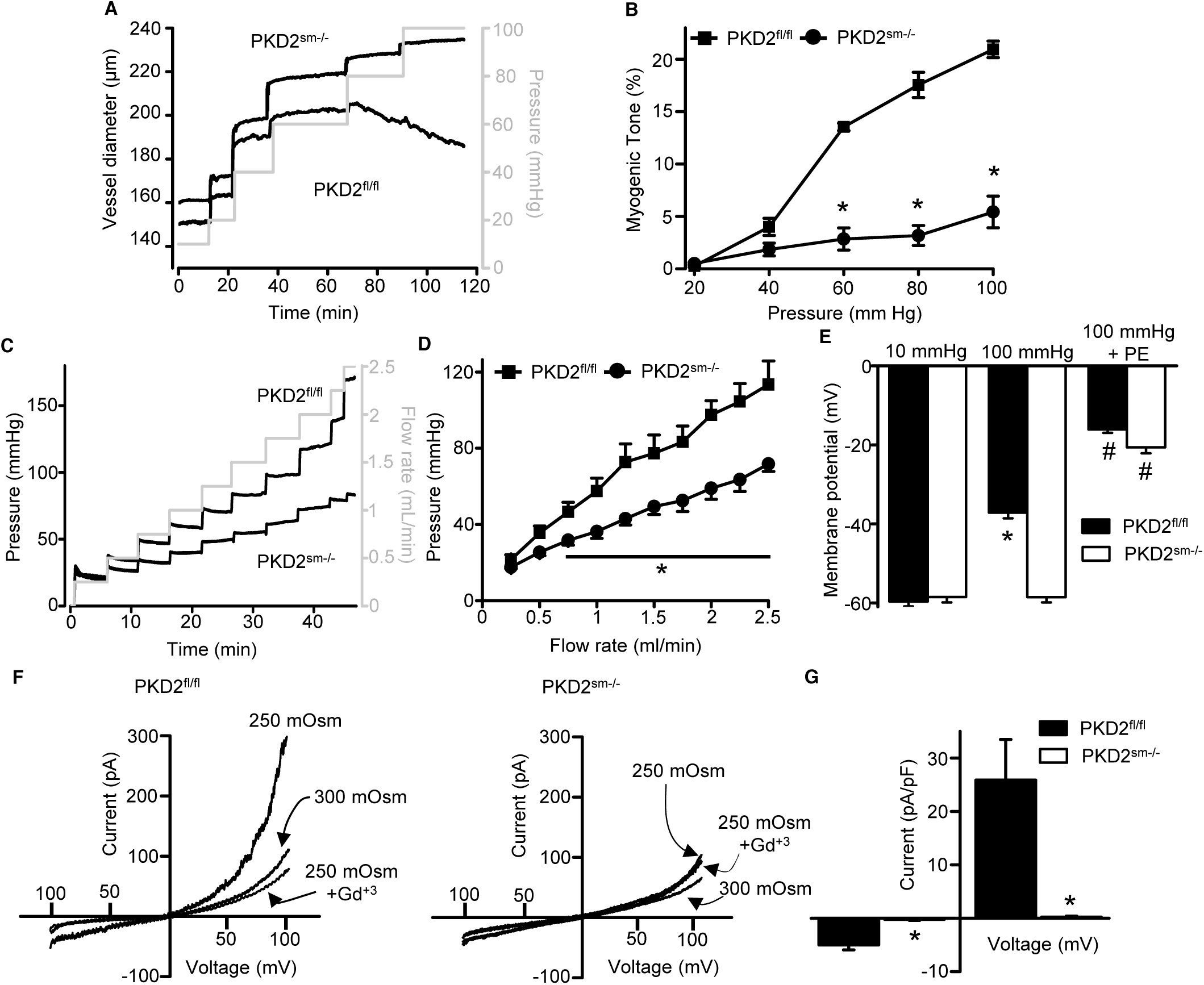
Pressure-induced vasoconstriction is reduced in PKD2^sm−/−^ hindlimb arteries. **A:** Representative traces illustrating diameter responses to intravascular pressure in gastrocnemius arteries of PKD2^fl/fl^ and PKD2^sm−/−^ mice. **B**: Mean data for myogenic tone in gastrocnemius arteries (PKD2^fl/fl^, n=5; PKD2^sm−/−^, n=6). **C**: Representative traces illustrating hindlimb perfusion pressure in response to increasing flow. **D**: Mean data for hindlimb perfusion pressure (PKD2^fl/fl^, n=6; PKD2^sm−/−^, n=4). **E:** Mean data for membrane potential recordings of pressurized hindlimb arteries in the absence or presence of phenylephrine (PE 1 μmol/L; PKD2^fl/fl^: 10 mmHg, n=11; 100 mmHg, n=10; 100 mmHg+PE, n=13 and PKD2^sm−/−^: 10 mmHg, n=11; 100 mmHg, n=10; 100 mmHg+PE, n=14). * indicates p<0.05 vs 10 mmHg in PKD2^fl/fl^. # indicates p<0.05 vs 100 mmHg in the same genotype. **F**: Representative voltage ramps illustrating I_Cat_ recorded in isotonic (300 mOsm) and hypotonic (250 mOsm) bath solution with and without Gd^3+^ (100 μmol/L) in the same PKD2^fl/fl^ and PKD2^sm−/−^ mouse hindlimb artery myocytes. **G**: Mean data for Gd^3^+-sensitive I_Cat_ recorded in hypotonic solution in PKD2^fl/fl^ and PKD2^sm−/−^ myocytes (n=5 for each). * indicates p<0.05 vs PKD2^fl/fl^.

A perfused hindlimb preparation was used to study vascular autoregulation in intact skeletal muscle. Increasing intravascular flow produced lower pressures in hindlimbs of PKD2^sm−/−^ mice than in those of PKD2^fl/fl^ mice (Figures 3C, D). For example, at a flow rate of at 2.5 ml/min the mean pressure in hindlimbs of PKD2^sm−/−^ mice were ~63.2 % of that in PKD2^fl/fl^ (Figures 3C, D). Sympathetic control of blood pressure occurs in part due to the activation of α_1_-adrenergic receptors in arterial myocytes. To investigate the contribution of myocyte PKD2 channels to α_1_-adrenergic receptor-mediated vasoconstriction, responses to phenylephrine, a selective α_1_-adrenergic receptor agonist, were measured. At constant flow, phenylephrine similarly increased pressure in PKD2^sm−/−^ and PKD2^fl/fl^ mouse hindlimbs (Supplemental Figure 5C).

### Pressure-induced membrane depolarization requires myocyte PKD2 channels in hindlimb arteries

To investigate the mechanism(s) by which smooth muscle cell PKD2 channels control contractility, membrane potential was measured in pressurized hindlimb arteries using glass microelectrodes. At 10 mmHg, the mean membrane potential of PKD2^fl/fl^ and PKD2^sm−/−^ arteries were similar at ~ −59.6 and −58.5 mV, respectively (Figure 3E, Supplemental Figure 5E). Increasing intravascular pressure to 100 mmHg depolarized PKD2^fl/fl^ arteries by ~22.5 mV, but did not alter the membrane potential of arteries from PKD2^sm−/−^ mice (Figure 3E, Supplemental Figure 5E). In contrast, phenylephrine depolarized arteries to similar membrane potentials in both genotypes (Figure 3E, Supplemental Figure 5E). These data suggest that pressure activates PKD2 channels in smooth muscle of hindlimb arteries, leading to depolarization and vasoconstriction.

### Swelling activates PKD2 channels in hindlimb artery myocytes

The contribution of PKD2 channels to mechanosensitive currents was investigated in hindlimb artery myocytes. Recent evidence indicates that recombinant PKD2 are voltage-dependent, generate outwardly rectifying currents and are primarily a Na^+^-permeant channel under physiological membrane potentials and ionic gradients ^7, 16^. Whole cell I_Cat_ was recorded using the whole-cell patch-clamp configuration with symmetrical Na^+^ solutions which had a reversal potential for Na^+^ (E_Na_) of 0 mV. In isotonic bath solution, at −100 mV mean I_Cat_ densities were similar in PKD2^fl/fl^ and PKD2^sm−/−^ myocytes (P<0.05, Figures 3F, G). In contrast, at +100 mV mean ICat density in PKD2^sm−/−^ myocytes (2.8±0.4 pA/pF) was 32.2% of that in PKD2^fl/fl^ myocytes (8.7±0.4 pA/pF, P<0.05, Figures 3F, G). These data are consistent with voltage-dependent activation of PKD2 channels in PKD2^fl/fl^ myocytes and the absence of this current in PKD2^sm−/−^ myocytes. Reducing bath solution osmolarity from 300 to 250 mOsm caused cell swelling and activated an outwardly rectifying I_Cat_ that increased 2.3-fold at −100 mV and 3.0-fold at +100 mV in myocytes of PKD2^fl/fl^ mice (Figure 3F, G). The swelling-activated I_Cat_ was inhibited by Gd^3+^, a non-selective cation channel blocker, in PKD2^fl/fl^ myocytes (Figure 3F, G). The mean reversal potential for I_Cat_ was the same in isoosmotic (−0.2±1.3 mV) and hypoosmotic solutions (0.1±3.8 mV; P>0.05). The reversal potential for I_Cat_ is the same as E_Na_, indicating that swelling activated a Na^+^ current (I_Na_). Reducing bath osmolarity did not activate a Gd^3+^-sensitive I_Cat_ in PKD2^sm–/–^ mouse hindlimb artery myocytes (Figure 3F, G). In contrast to the effects of cell swelling, phenylephrine activated similar amplitude I_Cat_s in hindlimb artery myocytes of PKD2^fl/fl^ and PKD2^sm–/–^ mice (Supplemental Figure 6). These data indicate that cell swelling, a mechanical stimulus, activates PKD2 channels, leading to a Na^+^ current in hindlimb artery myocytes.

### Myocyte PKD2 channels contribute to phenylephrine-induced vasoconstriction in mesenteric arteries

Mesenteric arteries were studied to determine whether myocyte PKD2 channels contribute to myogenic vasoconstriction in another arterial bed that is a major regulator of systemic blood pressure. In contrast to the loss of myogenic vasoconstriction in gastrocnemius arteries of PKD2^sm−/−^ mice, pressure- and depolarization-induced vasoconstriction were similar in mesenteric arteries of PKD2^sm−/−^ and PKD2^fl/fl^ mice (Supplemental Figures 7A, B, C). Myogenic tone was similar in PKD2^sm−/−^ and PKD2^fl/fl^ regardless of whether third, fourth, or fifth-order mesenteric arteries were studied (Supplemental Figure 7B). Passive diameter of mesenteric arteries was not altered by myocyte PKD2 knockout (Supplemental Figure 5A). Furthermore, the differential contribution of myocyte PKD2 channels to myogenic tone in gastrocnemius and mesenteric arteries was not due to size as passive diameters of first-order gastrocnemius and third-order mesenteric arteries were similar (Supplemental Figure 5A).

The splanchnic circulation receives considerable sympathetic innervation. To investigate the contribution of myocyte PKD2 channels to α_1_-adrenoceptor-mediated responses, phenylephrine-induced vasoconstriction was measured. Isometric contractions in mesenteric artery (first- and second-order) rings of PKD2^sm−/−^ mice were smaller than those in PKD2^fl/fl^ controls (Figures 4A, B). With 1 μmol/L phenylephrine, contractions in PKD2^sm−/−^ arteries were ~15.3 % of those in PKD2^fl/fl^ arteries, with maximal phenylephrine-induced contraction ~57.3 % of that in PKD2^fl/fl^ arteries (Figure 4B). The mean concentration of phenylephrine-induced half-maximal contraction (EC_50_, μmol/L) was slightly higher in PKD2^sm−/−^ (2.4±0.2) than PKD2^fl/fl^ (1.1±0.3) arteries (P<0.05, Figure 4B). In pressurized (80 mmHg) mesenteric (fourth- and fifth-order) arteries of PKD2^sm−/−^ mice, phenylephrine-induced vasoconstrictions were between ~50.9 and 64.8 % of those in PKD2^fl/fl^ controls (Figures 4C, D). Similar results were obtained with endothelium-denuded mesenteric arteries, indicating that attenuated vasoconstriction to phenylephrine was due to loss of PKD2 in myocytes (Supplemental Figure 7D, 7E). In contrast to attenuated phenylephrine-mediated vasoconstriction, angiotensin II-induced constriction was not different between PKD2^fl/fl^ and PKD2^sm−/−^ mesenteric arteries (Supplemental Figure 7F. These data indicate that arterial myocyte PKD2 channels are activated by distinct vasoconstrictor stimuli in arteries of different tissues, being essential for the myogenic response in hindlimb arteries and contributing to α_1_-adrenoceptor-mediated vasoconstriction in mesenteric arteries.

**Figure 4:**
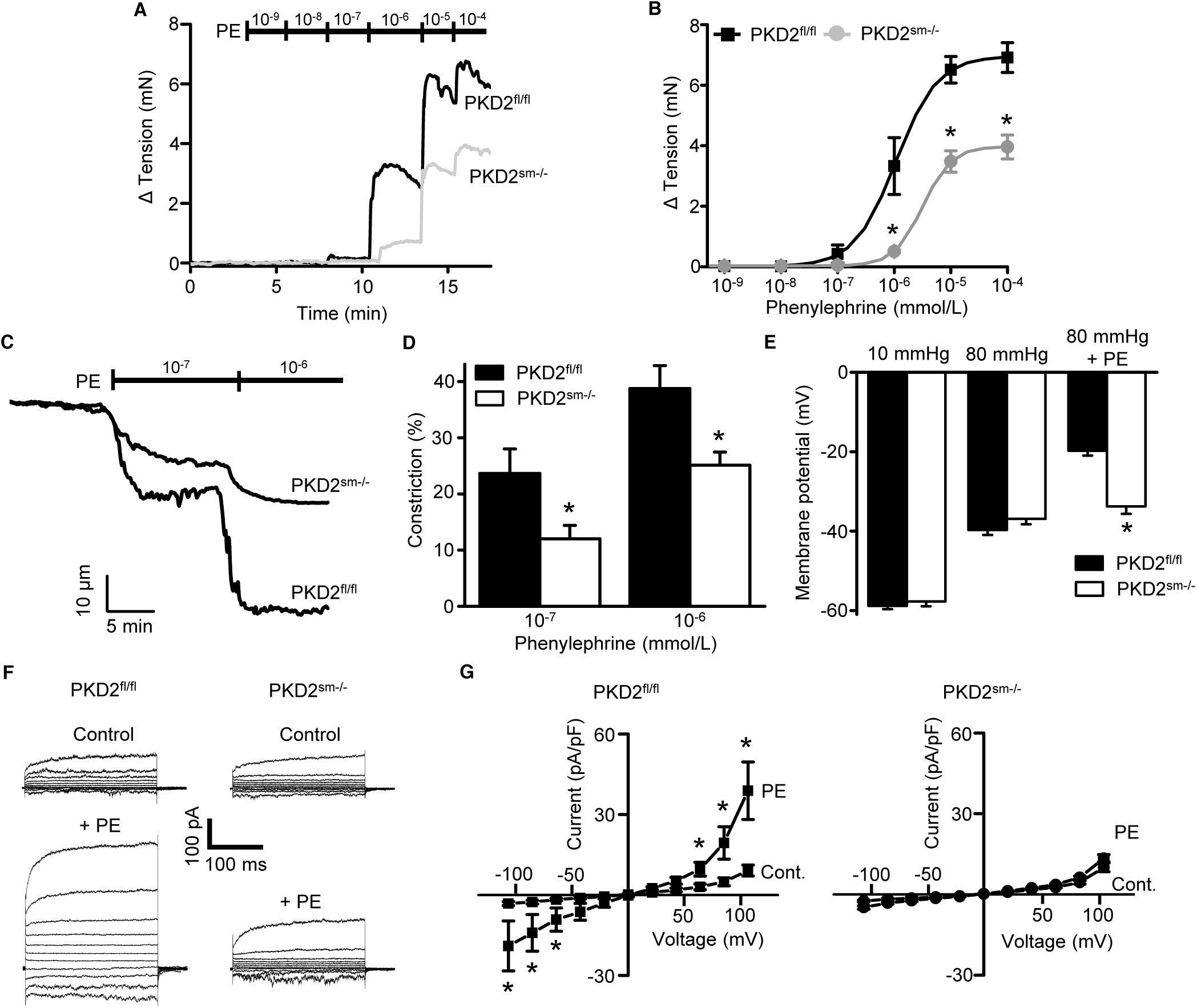
Phenylephrine-induced vasoconstriction is reduced in mesenteric arteries of PKD2^sm−/−^ mice. **A:** Original recordings of concentration-dependent, phenylephrine (PE)-induced contraction in mesenteric artery rings. **B**: Mean phenylephrine-induced contraction (PKD2^fl/fl^, n=5; PKD2^sm−/−^, n=6). **C**: Representative phenylephrine-induced vasoconstriction in pressurized fifth-order mesenteric arteries. **D**: Mean phenylephrine-induced vasoconstriction in pressurized fourth- and fifth-order mesenteric arteries (PKD2^fl/fl^ n=6 and PKD2^sm−/−^ n=6). **E:** Mean data for membrane potential recordings of pressurized mesenteric arteries in the absence or presence of phenylephrine (PE 1 μmol/L; PKD2^fl/fl^: 10 mmHg, n=13; 80 mmHg, n=9; 80 mmHg+PE, n=15 and PKD2^sm−/−^: 10 mmHg, n=11; 80 mmHg, n=12; 80 mmHg+PE, n=11). **F**: Original current recordings between −100 and +100 mV from a holding potential of −40 mV in PKD2^fl/fl^ and PKD2^sm−/−^ myocytes in control (Cont.) and PE (10 μmol/L). **G**: Mean paired data (PKD2^fl/fl^, n=5; PKD2^sm−/−^, n=4). * indicates p<0.05 vs control.

### α_1_-adrenergic receptors stimulate myocyte PKD2 channels, leading to membrane depolarization in mesenteric arteries

At an intravascular pressure of 10 mmHg, the membrane potential of PKD2^fl/fl^ and PKD2^sm−/−^ mesenteric arteries was similar (Figure 4E). An increase in pressure to 80 mmHg stimulated a similar depolarization in mesenteric arteries of PKD2^fl/fl^ and PKD2^sm−/−^ mice, to ~39.7 and 36.7 mV, respectively (Figures 4E, Supplemental Figure 7G). Phenylephrine application further depolarized PKD2^fl/fl^ mesenteric arteries by ~19.9 mV, but did not change the membrane potential of PKD2^sm−/−^ arteries (Figures 4E, Supplemental Figure 7G).

To examine phenylephrine-regulation of I_Cat_ in isolated mesenteric artery myocytes, the whole-cell patch-clamp configuration was used with symmetrical Na^+^ as the charge carrier. Basal I_Cat_ density was similar in mesenteric artery myocytes of PKD2^fl/fl^ and PKD2^sm−/−^ mice (Figure 4F, G). Phenylephrine activated outwardly rectifying I_Cat_, increasing density ~5.9- and 4.3-fold at −100 mV and +100 mV, respectively in PKD2^fl/fl^ cells (Figure 4F, G). In contrast, phenylephrine did not alter ICat in PKD2^sm−/−^ myocytes (Figure 4F, G). The mean reversal potential for ICat in PKD2^fl/fl^ myocytes was the same in control (0.2±1 mV) and phenylephrine (1.3±0.7 mV; P>0.05) and is similar to E_Na_. These data indicate that phenylephrine activates PKD2 channels, leading to a Na^+^ current (I_Na_) in mesenteric artery myocytes. In contrast to the differential effects of phenylephrine, swelling activated similar amplitude ICat in mesenteric artery myocytes of PKD2^fl/fl^ and PKD2^sm−/−^ mice (Supplemental Figure 8). These data indicate that phenylephrine, but not cell swelling, activates PKD2 channels in mesenteric artery myocytes.

### PKD2 channel knockout in arterial myocytes attenuates hypertension

We tested the hypothesis that arterial myocyte PKD2 channels are associated with the increase in blood pressure during hypertension. Angiotensin II-infusion is a well-established method to produce a stable elevation in mean arterial pressure. Blood pressure was measured following implantation of subcutaneous osmotic minipumps that infused angiotensin II or saline in PKD2^sm−/−^ and PKD2^fl/fl^ mice. Angiotensin II steadily increased MAP to a plateau of ~155.6 mmHg in PKD2^fl/fl^ mice and to ~134.6 mmHg in PKD2^sm−/−^ mice (Figure 5A). The angiotensin II-induced increase in MAP (ΔMAP) was ~25.6 % smaller in PKD2^sm−/−^ than PKD2^fl/fl^ mice (Figure 5A). Saline infusion did not alter blood pressure in either PKD2^sm−/−^ or PKD2^fl/fl^ mice. These data indicate that myocyte PKD2 channel knockout reduces both the absolute systemic blood pressure and the increase in blood pressure during hypertension.

**Figure 5:**
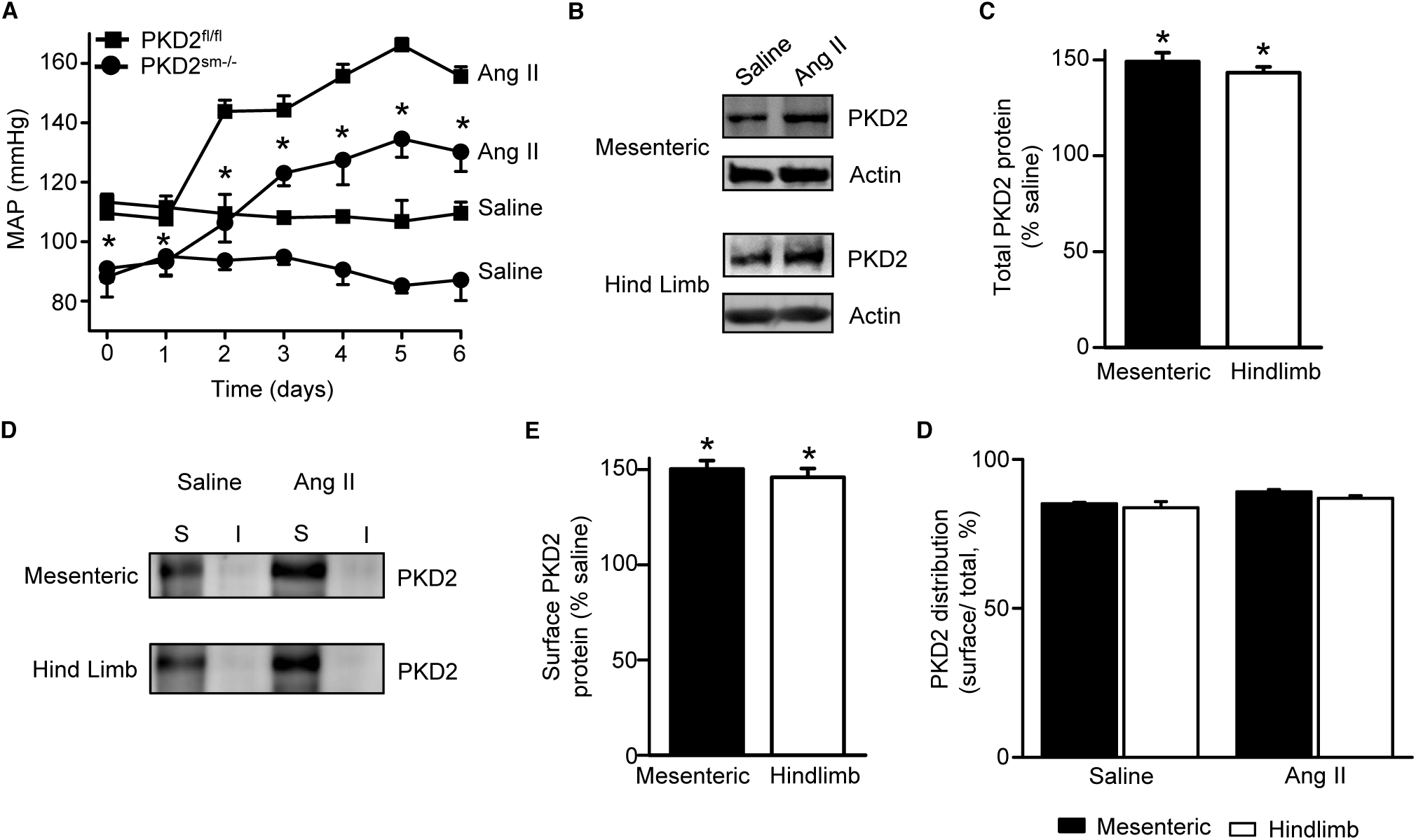
Angiotensin II-induced hypertension is attenuated in PKD2^sm−/−^ mice. **A:** Telemetric blood pressure time course showing the development of angiotensin II-induced hypertension in PKD2^fl/fl^ (n=6) and PKD2^sm−/−^ mice (n=9). Osmotic minipumps containing either saline or angiotensin II were implanted one day prior to day 0. * indicates p<0.05 ang II PKD2^sm−/−^ vs ang II PKD2^fl/fl^. **B**: Western blots illustrating total PKD2 protein in mesenteric and hindlimb arteries of saline- and angiotensin II-treated control mice. **C**: Mean total PKD2 protein in mesenteric and hindlimb arteries of angiotensin II-treated mice compared to saline control (n=8 for each group). * indicates p<0.05 vs saline in the same arterial preparation. **D**: Western blots showing surface and intracellular PKD2 protein in arteries of saline- and angiotensin II-treated mice. **E**: Mean surface PKD2 protein in mesenteric and hindlimb arteries of angiotensin II-treated mice compared to saline control (n=8 for each group). * indicates p<0.05 vs saline in the same arterial preparation. **F**: Mean data illustrating the percentage of total PKD2 located at the surface in mesenteric and hindlimb arteries of saline- and angiotensin II-treated mice (n=8 for each group).

### Hypertension is associated with an upregulation of both total and surface PKD2 proteins in systemic arteries

To investigate mechanisms by which arterial myocyte PKD2 channels may be associated with an increase in blood pressure during hypertension, total and surface proteins were measured in arteries. PKD2 total protein in mesenteric and hindlimb arteries of angiotensin II-induced hypertensive mice were ~149.2 and 143.4 %, respectively, of those in normotensive mice (Figures 5B, C). Arterial biotinylation revealed that surface PKD2 protein in mesenteric and hindlimb arteries of hypertensive mice were also ~150.3 and 145.9 % of those in controls (Figure 5D, E). In contrast, cellular distribution of PKD2 was similar in mesenteric and hindlimb arteries of normotensive and hypertensive mice, with ~85 % of protein located in the plasma membrane (Figures 5D). These data indicate that during hypertension, an increase in total PKD2 protein leads to an increase in the abundance of surface PKD2 channels.

### Myocyte-specific PKD2 channel knockout induces vasodilation and prevents arterial remodeling during hypertension

To test the hypothesis that the reduction in systemic blood pressure in PKD2^sm−/−^ mice during hypertension was due to vasodilation, the contractility of pressurized (80 mmHg) mesenteric arteries was measured using myography. Phenylephrine-induced vasoconstrictions in angiotensin II-treated PKD2^sm−/−^ mouse arteries were between ~67.6 and 71.1 % of those in PKD2^fl/fl^ arteries (Figure 6A). In contrast, myogenic tone was similar in arteries of angiotensin II-treated PKD2^sm−/−^ and PKD2^fl/fl^ mice (Figure 6B). These results indicate that knockout of arterial myocyte PKD2 channels attenuates phenylephrine-induced vasoconstriction during hypertension. These data also suggest that hypertension does not promote the emergence of a mechanism by which PKD2 channels contribute to the myogenic response in mesenteric arteries.

**Figure 6:**
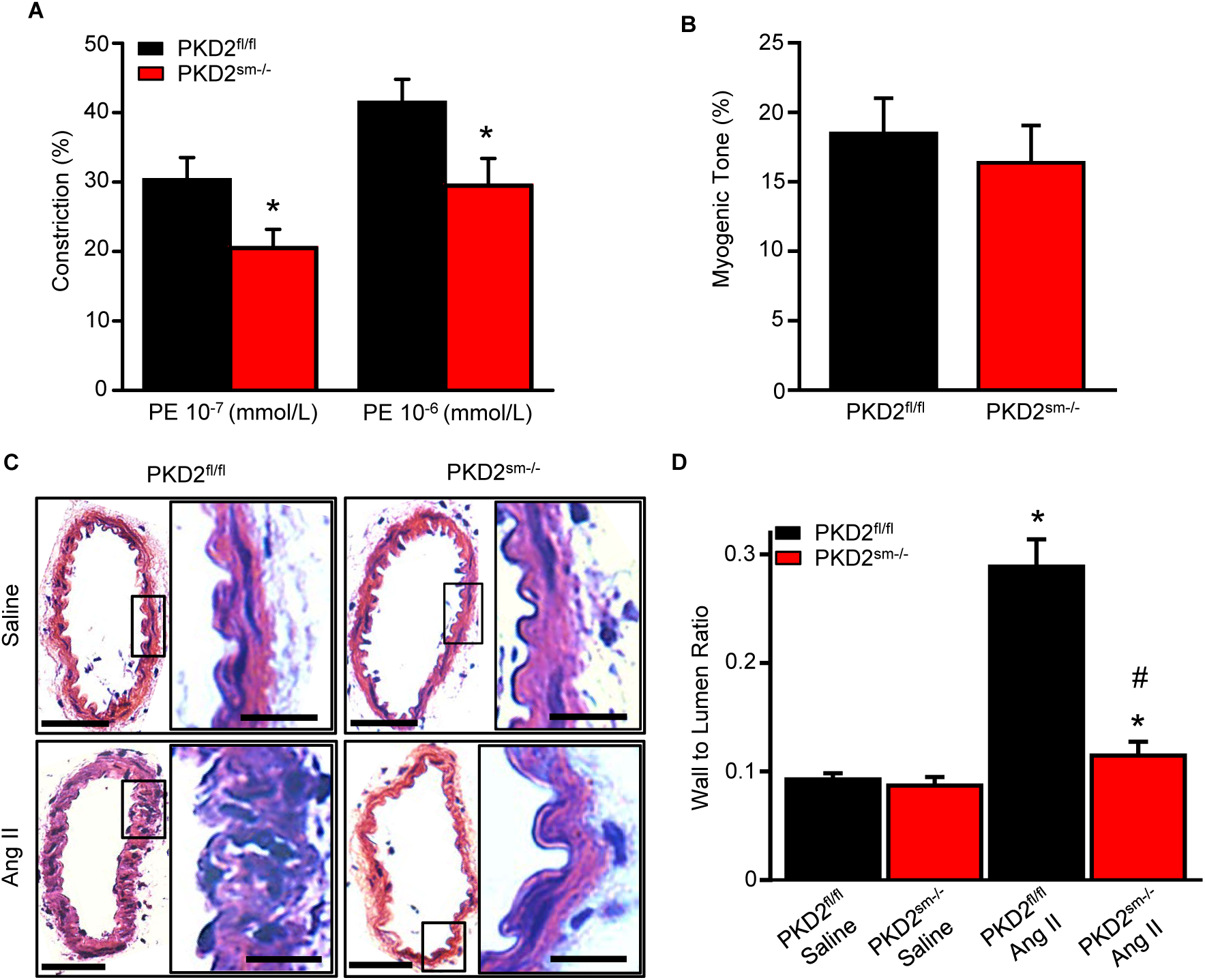
Arterial myocyte PKD2 knockout attenuates vasoconstriction and arterial wall remodeling during hypertension. **A:** Mean phenylephrine-induced vasoconstriction in pressurized (80 mmHg) mesenteric arteries from angiotensin II-treated mice (PKD2^fl/fl^, n=8; PKD2^sm−/−^, n=8). **B**: Mean data of myogenic tone at 80 mmHg in mesenteric arteries from Ang II treated mice (PKD2^fl/fl^, n=8; PKD2^sm−/−^, n=8). **C**: Representative H&E stained sections of fourth-order mesenteric arteries from saline and angiotensin II-treated PKD2^fl/fl^ and PKD2^sm−/−^ mice with insets. Low- and high-magnification scale bars are 50 and 25 μm, respectively. **D**: Mean data for wall-to-lumen ratio (PKD2^fl/fl^, n=17; PKD2^sm−/−^, n=20; PKD2^fl/f^ ang II, n=23; PKD2^sm−/−^ ang II, n=22). * indicates p<0.05 vs saline in the same mouse model and # is ang II PKD2^sm−/−^ vs ang II PKD2^fl/fl^.

Hypertension is associated with inward remodeling of vasculature ^17^. To investigate the involvement of myocyte PKD2 channels in pathological remodeling, arterial sections from angiotensin II-treated PKD2^sm−/−^ and PKD2^fl/fl^ mice were imaged and analyzed. In saline-treated PKD2^sm−/−^ and PKD2^fl/fl^ mice, the wall-to-lumen ratio of mesenteric arteries was similar (Figures 6C, D). Angiotensin II infusion increased the artery wall-to-lumen ratio ~2.9-fold in PKD2^fl/fl^ mice, but only 1.3-fold in PKD2^sm−/−^ mice, or ~89.6 % less (Figures 6C, D). These data suggest that PKD2 knockout in myocytes attenuates arterial remodeling during hypertension.

## Discussion

Previous studies performed *in vitro* have suggested that arterial myocytes express several different TRP channels, but *in vivo* physiological functions of these proteins are unclear. Here, we generated the first inducible, smooth muscle-specific knockout of a TRP channel, PKD2, to investigate blood pressure regulation by this protein. We show that tamoxifen-induced, smooth muscle-specific PKD2 knockout dilates resistance-size systemic arteries and reduces blood pressure. Data indicate that heterogeneous vasoconstrictor stimuli activate PKD2 channels in arterial myocytes of different tissues. PKD2 channel activation leads to a I_Na_ in myocytes, which induces membrane depolarization and vasoconstriction. Furthermore, we show that hypertension is associated with an increase in plasma membrane-resident PKD2 channels. PKD2 channel knockout in myocytes of hypertensive mice caused vasodilation, prevented arterial remodeling and lowered systemic blood pressure. In summary, using a novel inducible, conditional knockout model we show that arterial myocyte PKD2 channels are active *in vivo*, which increases physiological systemic blood pressure. We also show that arterial myocyte PKD2 channels are upregulated during hypertension and genetic knockout reduces high blood pressure.

The regulation of blood pressure by arterial myocyte TRP proteins and involvement in hypertension are poorly understood ^2^. This lack of knowledge is due to several primary factors. First, it is unclear which TRP channel family members are expressed and functional in myocytes of resistance-size systemic arteries that control blood pressure. The identification of a TRP channel in blood vessels that do not regulate systemic blood pressure, such as aorta and cerebral arteries, does not necessarily indicate that the same protein is present in arteries that do control blood pressure ^2^. Second, TRP channel expression has often been reported in myocytes that had undergone cell culture or in whole vasculature that contains many different cell types. These studies create uncertainty as to which TRP proteins are present specifically in contractile myocytes of resistance-size systemic arteries ^2^. Third, the lack of specific modulators of individual TRP channel subfamily members has significantly hampered studies aimed at identifying *in vitro* and *in vivo* functions of these proteins. Fourth, TRP channels are expressed in many different cell types other than arterial myocytes. Blood pressure measurements in global, non-inducible TRP knockout mice generated unexpected and/or contrasting results to those anticipated. Myocyte TRPM4 channel activation leads to vasoconstriction in isolated cerebral arteries, but global, constitutive knockout of TRPM4 elevated blood pressure in mice ^3, 18^. Arterial myocyte TRPC6 channel activation stimulates cerebral artery constriction, but global, constitutive TRPC6 knockout caused vasoconstriction and elevated blood pressure in mice ^4, 19^. TRPV4 channel activation in arterial myocytes leads to vasodilation in isolated vessels, but blood pressure in TRPV4^−/−^ mice was the same or lower than controls^5, 6^. Alternative approaches, such as the inducible, cell-specific knockout model developed here, are valuable to advance knowledge of blood pressure regulation by arterial myocyte TRP channels.

Previous studies performed *in vitro* that investigated vasoregulation by PKD2 channels produced a variety of different findings. RNAi-mediated knockdown of PKD2 inhibited swelling-activated non-selective cation currents (I_Cat_) in rat cerebral artery myocytes and reduced myogenic tone in pressurized cerebral arteries ^10^. In contrast, PKD2 siRNA did not alter myogenic tone in mesenteric arteries of wild-type mice, but PKD siRNA increased both stretch-activated cation currents and myogenic vasoconstriction in arteries of constitutive, myocyte PKD1 knockout mice ^11^. From these results, the authors suggested that myocyte PKD2 channels inhibit mesenteric vasoconstriction ^11^. Global knockout of PKD2 is embryonic-lethal in mice due to cardiovascular and renal defects, prohibiting study of the loss of protein in all tissues on vascular function ^20, 21^. In global PKD2^+/−^ mice, an increase in actin and myosin expression lead to larger phenylephrine-induced contractions in aorta and mesenteric arteries ^12, 13^. Here, using an inducible cell-specific knockout model, we show that swelling, a mechanical stimulus, activates PKD2 currents in myocytes of skeletal muscle arteries, whereas phenylephrine activates PKD2 currents in myocytes of mesenteric arteries. Regardless of the stimulus involved, PKD2 activation leads to membrane depolarization and vasoconstriction in both artery types. We demonstrate for the first time that PKD2 channels are present in arterial myocytes of skeletal muscle arteries. PKD2 appears to be the second TRP protein to have been described in this cell type, TRPV1 being the other ^22^.

Few studies have measured plasma membrane currents through either recombinant PKD2 channels or PKD2 channels expressed in native cell types. PKD2 channels have been observed at the cell surface, with evidence suggesting that PKD1 is required for PKD2 translocation, although the ability of PKD1 to perform this function has been questioned ^7 23^. Here, using arterial biotinylation we show that 85 % of PKD2 protein is located in the plasma membrane in both mesenteric and hindlimb arteries, consistent with our previous data in cerebral arteries ^10^. PKD2 ion permeability has also been a matter of debate. Recombinant PKD2 was initially shown to both conduct Ca^2+^ and to be inhibited by Ca^2+^ ^23, 24^. To increase surface-trafficking of recombinant PKD2, a recent study generated a chimera by replacing the pore of PKD2-L1, a related channel that readily traffics to the plasma membrane, with the PKD2 channel pore ^7^. This channel, which contained the PKD2 channel filter, generated outwardly-rectifying whole cell currents and was more selective for Na^+^ and K^+^ than Ca^2+^, with Ca^2+^ only able to permeate in an inward direction ^7^. We also show that PKD2 channels in hindlimb and mesenteric artery myocytes are outwardly rectifying, permeant to Na^+^, as this was by far the predominant cation present in solutions, and currents reversed at the predicted E_Na_ of 0 mV. At physiological arterial potentials of ~-60 to −40 mV, PKD2 channel opening would produce an inward Na^+^ current, leading to myocyte membrane depolarization and voltage-dependent Ca^2+^ (Ca_v_1.2) channel activation ^25^. The ensuing increase in intracellular Ca^2+^ concentration would stimulate vasoconstriction and an elevation in blood pressure, as observed here. PKD2 channel knockout abolishes the stimulus-induced inward Na^+^ current, attenuating vasoconstriction.

The splanchnic and skeletal muscle circulations account for up to 35 and 80 (during physical exertion) % of cardiac output, respectively. Changes in arterial contractility within these organs leads to significant modification of total peripheral resistance and systemic blood pressure. We demonstrate that myocyte PKD2 channels regulate contractility in both of these physiologically important circulations. Furthermore, our data demonstrate that distinct mechanical and chemical stimuli activate PKD2 channels in these arteries to produce vasoconstriction. Intravascular pressure stimulates PKD2 channels in myocytes of hindlimb arteries, leading to vasoconstriction, whereas myocyte PKD2 channels contribute to phenylephrine-induced vasoconstriction in mesenteric arteries. Mice lacking α1-adrenergic receptors are hypotensive and have reduced vasoconstrictor responses to phenylephrine in mesenteric arteries, highlighting the relevance of this pathway to blood pressure regulation ^26^. Surprisingly, we found that myocyte PKD2 channels do not contribute to the myogenic response in mesenteric arteries or to phenylephrine-induced vasoconstriction in hindlimb arteries. This differential regulation of contractility by myocyte PKD2 channels is not due to arterial size. These results show that PKD2 is functional in arterial myocytes of different organs, but stimuli that activate these channels are vascular bed-specific. In contrast to responses to intravascular pressure and phenylephrine, angiotensin II-induced vasoconstriction did not require myocyte PKD2 channels in either hindlimb or mesenteric arteries. These data indicate that myocyte PKD2 knockout did not alter responses to all vasoactive agents and support the conclusion that myocyte PKD2 channels are activated by specific stimuli in the hindlimb and mesenteric circulations. This finding strengthens the concept that the regulation and function of a protein is not homogeneous in the circulation and findings in one arterial bed should not simply be translated to other vasculature.

It was outside of the central aim of this study and beyond scope to determine the differential mechanisms by which pressure and adrenergic receptors activate PKD2 channels in arteries of different organs. Pressure-induced vasoconstriction was described over 100 years ago and yet the mechanosensor, intracellular signal transduction pathways and the ion channels that produce this physiological response are still unresolved ^27^. Candidates for the pressure mechanosensor include, one or more proteins that have been proposed in smooth muscle or non-smooth muscle cell types, proteins that have not yet been discovered, elements of the cytoskeleton, or ion channels that act alone, in series or in parallel with other proteins and may be homomultimeric or heteromultimeric proteins. Importantly, we show here that there is not a singular, homogeneous mechanism that mediates the myogenic response in all artery types, adding additional complexity to the derivation of this mechanosensing signaling event. It is possible that the intravascular pressure mechanosensors are different in hindlimb and mesenteric artery myocytes. If this is the case, the mechanosensor protein(s) present in hindlimb artery myocytes activates PKD2 channels, whereas the mechanosensor(s) in mesenteric artery myocytes does not stimulate PKD2. The intracellular signaling pathways by which pressure and α_1_-adrenergic receptors activate PKD2 channels may also not be the same or may couple differently in myocytes of each artery type. Intracellular pathways that transduce the pressure signal to activate ion channels are also not resolved, although G_q11_-coupled phospholipase C (PLC) activation is one candidate ^28^. α_1_-adrenergic receptors also activate G_q/11_, which based on data here appears to couple to PKD2 channels in myocytes of mesenteric, but not hindlimb, arteries. If pressure and α_1_-adrenergic receptors both stimulate G_q11_, local or global proximity signaling to PKD2 channels may underlie differential signaling in each artery type. When considering the molecular identity of the channels that are involved, PKD2 can form both homotetramers and heteromultimers with other channels, including TRPC1 and TRPV4 ^23, 29–32^. Whether PKD2 forms homomers or heteromers with other TRP channels in arterial myocytes is unclear, but differential PKD2 channel tetramerization may also underlie the distinct activation of these proteins by pressure and α1-adrenergic receptors in hindlimb and mesenteric arteries. Other channels, including TRPC6, TRPM4 and ANO1 contribute to myogenic tone in cerebral arteries, but their relationship to PKD2-mediated responses in mesenteric and hindlimb arteries is unclear ^33–35^. Conceivably, other ion channels may signal in series to PKD2, with the proteins involved and the sequence of events differing in mesenteric and hindlimb artery myocytes. Given the large number of unresolved signaling elements and the possible permutations that would require study, it was beyond scope to determine the differential mechanisms by which pressure and adrenergic receptors activate PKD2 channels in arteries of different organs. Future studies should investigate these possibilities.

Having established that myocyte PKD2 channels regulate arterial contractility and systemic blood pressure, we investigated whether hypertension is associated with an alteration in PKD2 expression, surface protein and function. Total and surface PKD2 protein were both higher in mesenteric and hindlimb arteries of hypertensive mice than in normotensive controls, indicating upregulation. PKD2 protein was primarily located in the plasma membrane in arteries of normotensive and hypertensive mice demonstrating that the distribution of channels was unchanged. These data indicate that hypertension is associated with an increase in the amount of PKD2 protein, which traffics to the plasma membrane, thereby increasing surface channel number. Myocyte-specific PKD2 knockout reduced both phenylephrine-induced vasoconstriction and systemic blood pressure and prevented an increase in wall-to-lumen thickness in hypertensive mice. The angiotensin II-induced elevation in blood pressure was smaller in PKD2^sm−/−^ mice than in PKD2^fl/fl^ controls, supporting the concept that a higher abundance of surface PKD channels in arterial myocytes was associated with vasoconstriction. Our data demonstrate that myocyte PKD2 channels are upregulated during hypertension and that PKD2 targeting reduces vasoconstriction, blood pressure and arterial remodeling during hypertension, eliciting multi-modal benefits.

When considering our findings, a translation to human diseases that are associated with PKD2 protein is warranted. Autosomal Dominant Polycystic Kidney Disease (ADPKD) occurs due to mutations in either PKD1 or PKD2 and is the most prevalent monogenic human disease worldwide, affecting 1 in 1000 people ^36^. ADPKD is characterized by the appearance of renal cysts, although a significant proportion of patients with apparently normal renal function develop hypertension prior to the development of cysts ^36–38^. Currently, more than 275 human variants in PKD2 have been identified (Autosomal Dominant Polycystic Kidney Disease Mutation Database, Mayo Clinic; http://pkdb.pkdcure.org). Whether mutations in systemic artery myocyte PKD2 channels modify vascular contractility and blood pressure remains to be determined, but is rational to investigate given findings here. Future studies should investigate these possibilities.

In summary, we show that arterial myocyte PKD2 channels are activated by distinct stimuli in arteries of different tissues, increase systemic blood pressure, are upregulated during hypertension and genetic knockout *in vivo* leads to vasodilation and a reduction in both physiological and high blood pressure.

## Author Contributions

S.B., C.F-P, R.H., M.D.L., P.M., K.W.E., S.K.B, Q.W., K.P.K. and conducted experiments, acquired data, and analyzed data. S.B. and J. H. J. designed research studies and wrote the manuscript.

## Acknowledgements

This study was supported by NIH/NHLBI grants HL67061, 133256 and HL137745 to J.H.J, American Heart Association (AHA) Scientist Development Grants to S.B and M.D.M and AHA postdoctoral fellowship to R.H.

## SUPPLEMENTAL MATERIAL

**Supplemental Table 1.**
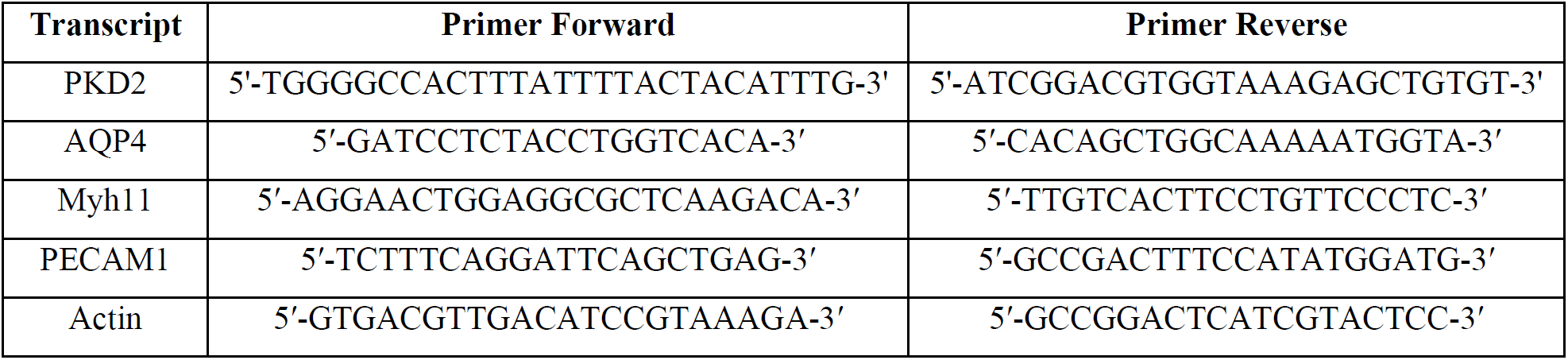
Primers used for RT PCR. The PKD2 forward primer recognizes nucleotides in exon 9 and 10 and the reverse primer was aligned with a sequence in exon 13.

## SUPPLEMENTAL FIGURE LEGENDS

**Supplemental Figure 1:**
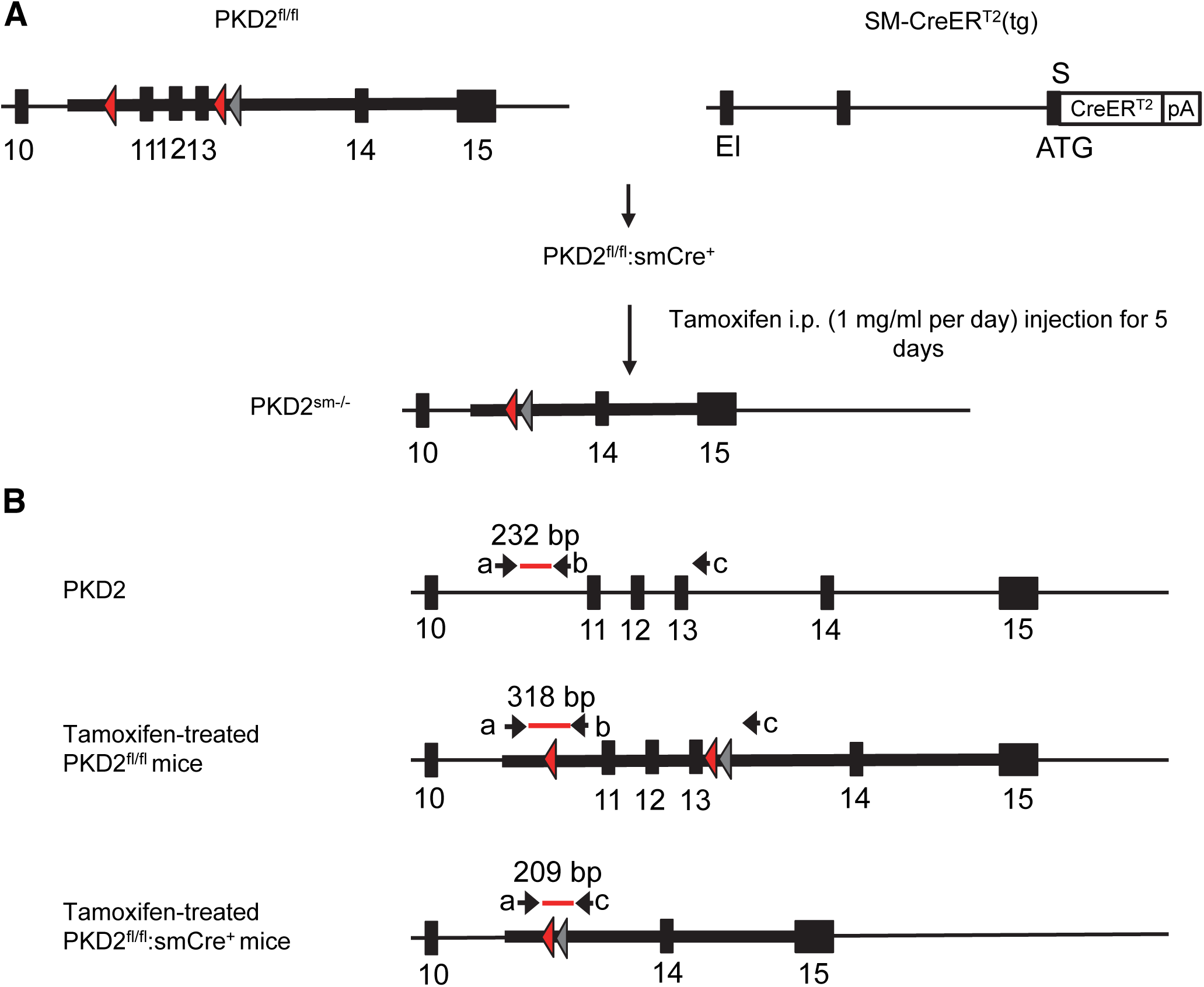
**A**: Schematic representation of the breeding protocol used to generate PKD2^fl/fl^:smCre^+^ mice and *Pkd2* recombination produced by tamoxifen activation of Cre recombinase. **B**: Genotyping of mice using PCR. Shown are the annealing locations of genomic PCR primers a, b, c and the size of the PCR product amplified. Red and gray triangles represent loxP and Frt sites, respectively.

**Supplemental Figure 2:**
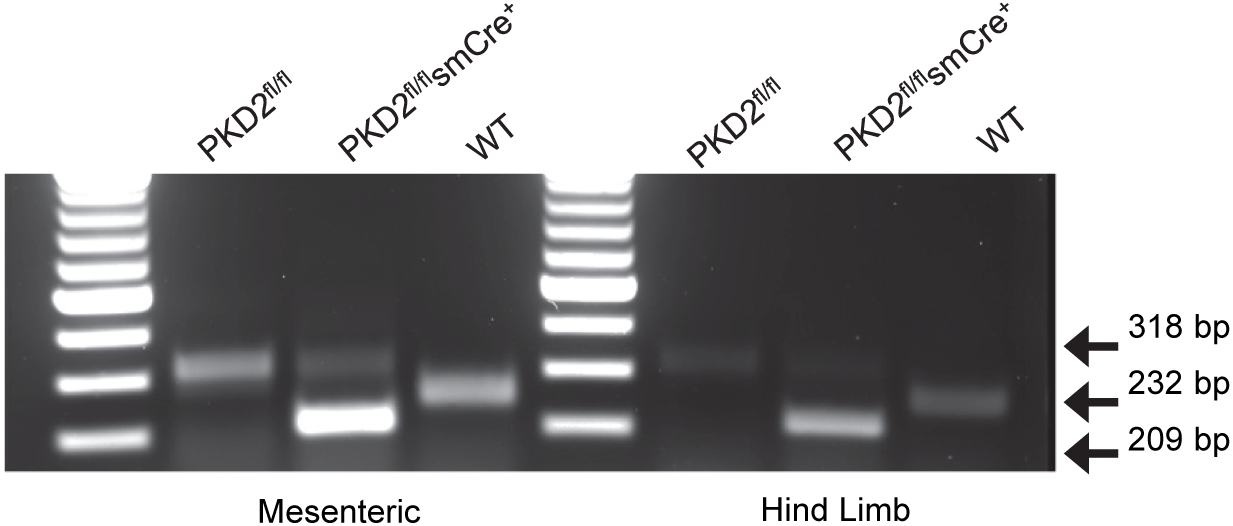
Genotyping of mouse lines. Ethidium bromide gel illustrating PCR products in vasculature of C57BL/6J mice and tamoxifen-treated PKD2^fl/fl^ and PKD2^fl/fl^:smCre^+^ mice.

**Supplemental Figure 3:**
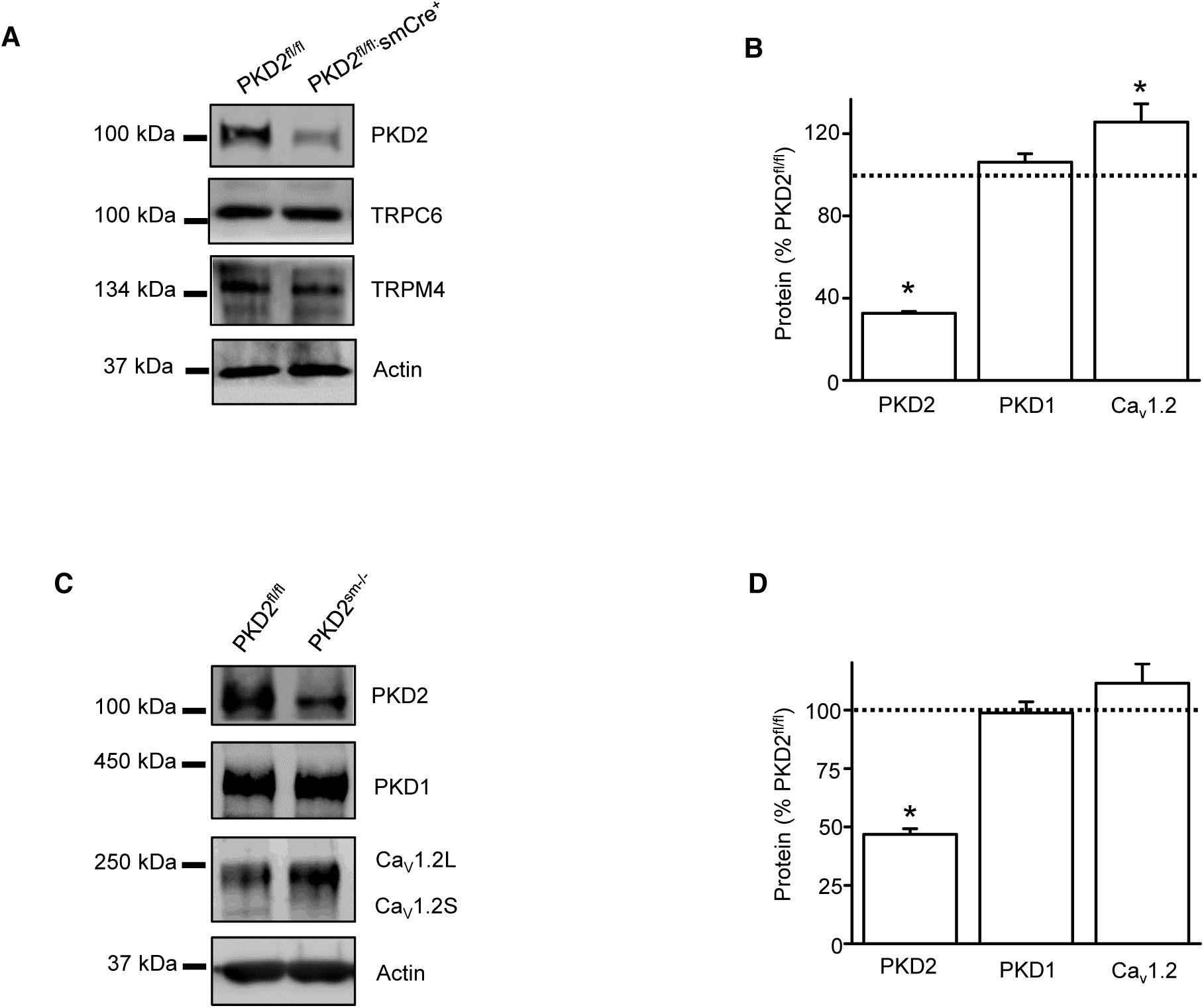
PKD2 protein is lower in aorta, mesenteric and hindlimb arteries from tamoxifen-treated PKD2^fl/fl^:smCre^+^ mice. **A:** Western blots illustrating PKD2 protein was lower in mesenteric arteries of tamoxifen-treated PKD2^fl/fl^:smCre^+^ mice, whereas other proteins were similar. **B**: Mean data for proteins in hindlimb arteries of PKD2^sm−/−^ mice (n=4–6). **C**: Western blots showing proteins in aorta. Ca_v_1.2L, full-length Ca_v_1.2; Ca_v_1.2S, short Ca_v_1.2. **D**: Mean data from aorta (n=4). * indicates p<0.05.

**Supplemental Figure 4:**
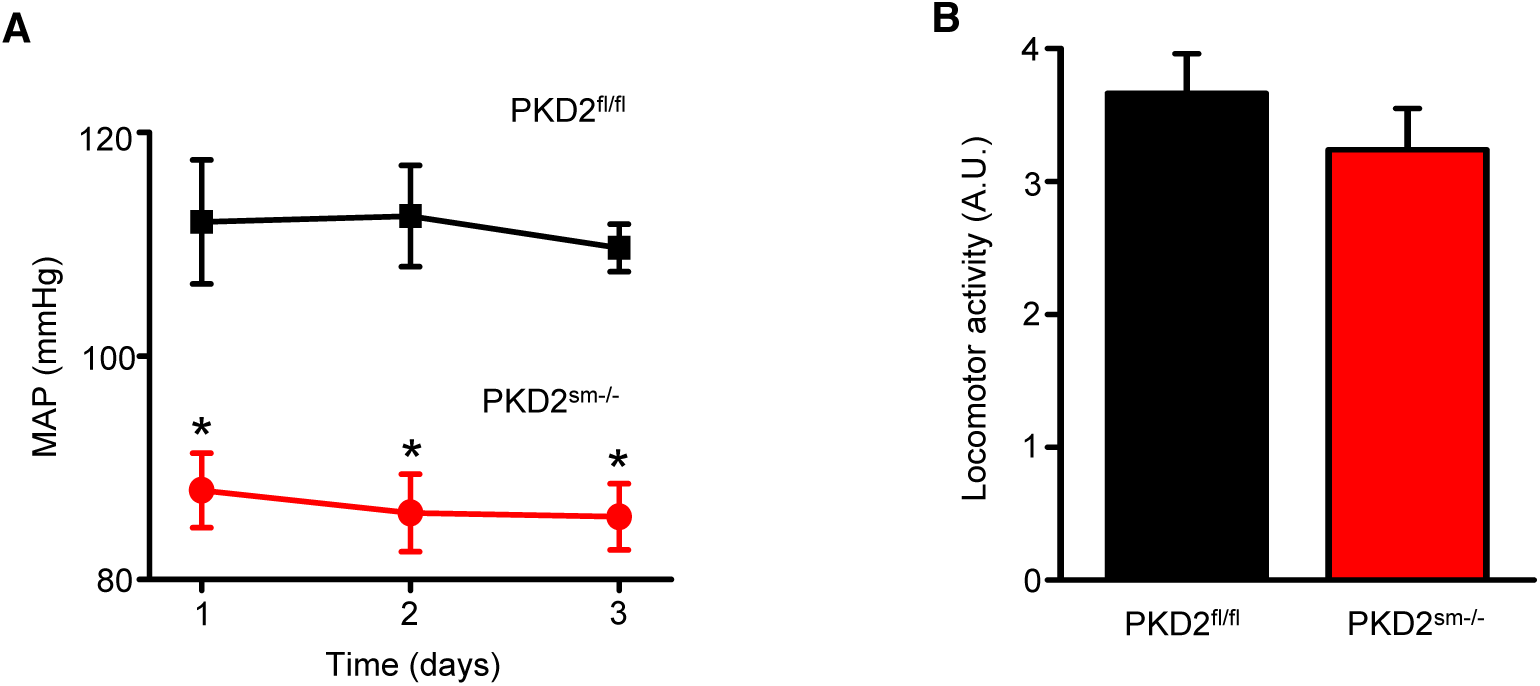
Lower blood pressure is sustained in PKD2^sm−/−^ mice. A: Mean arterial blood pressure (MAP) in PKD2^sm−/−^ and PKD2^fl/fl^ mice, n=6 per group. **B:** Mean data of locomotor activity, n=6 per group.

**Supplemental Figure 5:**
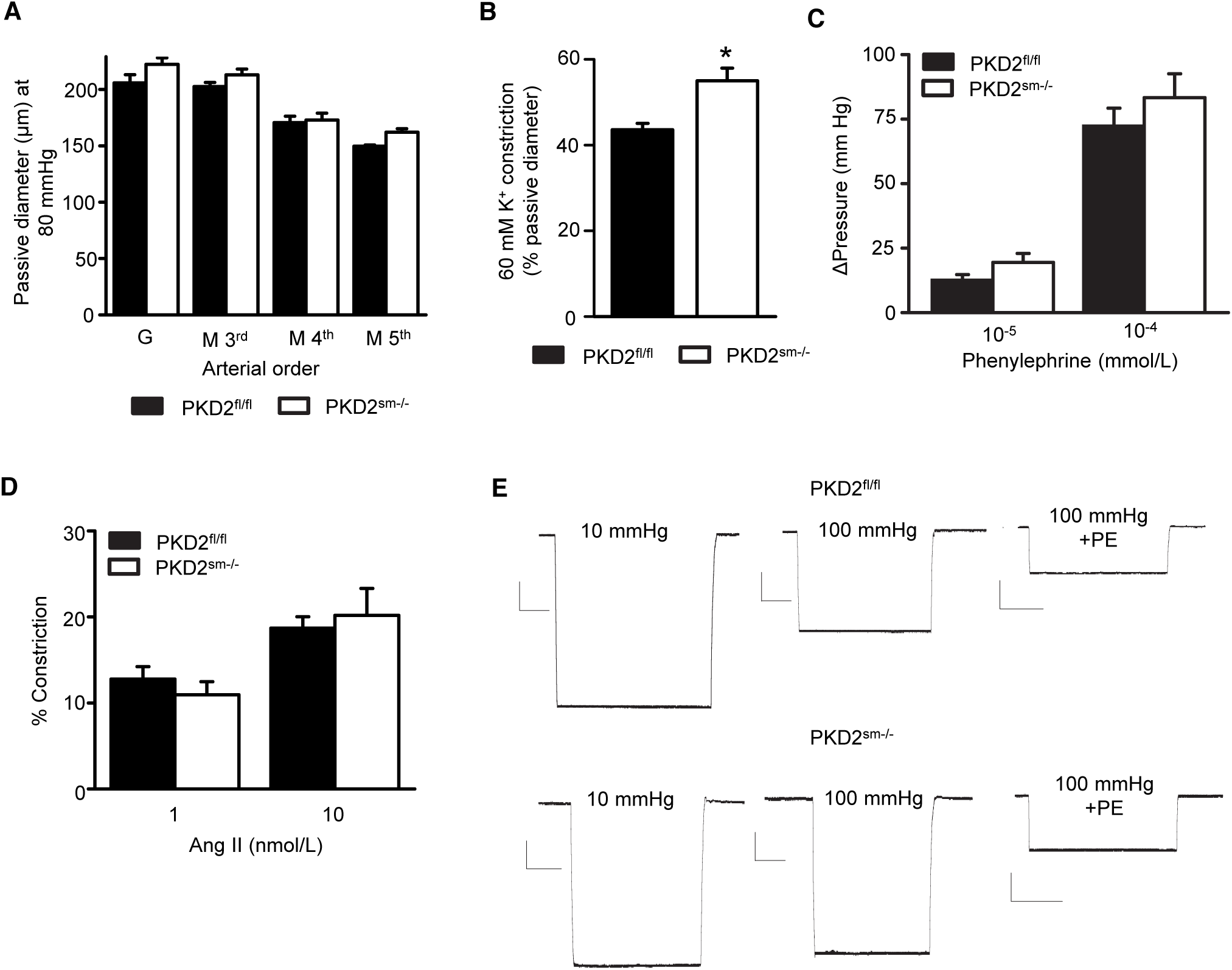
Myocyte PKD2 knockout attenuates pressure-induced membrane depolarization, but does not alter phenylephrine or angiotensin II-induced vasoconstriction in hindlimb arteries. **A:** Mean passive diameter at 80 mmHg of first-order gastrocnemius arterioles (G) and third-, fourth- and fifth-order mesenteric arteries (M) (PKD2^fl/fl^: G, n= 6; M3^rd^ n=4; M4^th^ n=5; M5^th^ n=4 and PKD2^sm−/−^: G, n= 6; M3^rd^ n=7; M4^th^ n=4; M5^th^ n=4). **B:** Mean data for 60 mmol/L K^+^-induced constriction in pressurized (100 mmHg) gastrocnemius arterioles from PKD2^fl/fl^ (n=4) and PKD2^sm−/−^ (n=5) mice. * indicates p<0.05. **C**: Mean data of phenylephrine-induced pressure responses in the intact hindlimb preparation (PKD2^fl/fl^, n=5 and PKD2^sm−/−^, n=4). **D**: Mean data of angiotensin II-induced constriction in hindlimb arteries pressurized to 100 mmHg (PKD2^fl/fl^, n=5 and PKD2^sm−/−^, n=6). **E**: Representative traces of microelectrode impalements under indicated conditions illustrating that pressure-induced depolarization is attenuated in gastrocnemius arteries of PKD2^sm−/−^ mice. Phenylephrine (PE) = 1 μmol/L. Scale bars: Y= 10 mV, X=20 s.

**Supplemental Figure 6:**
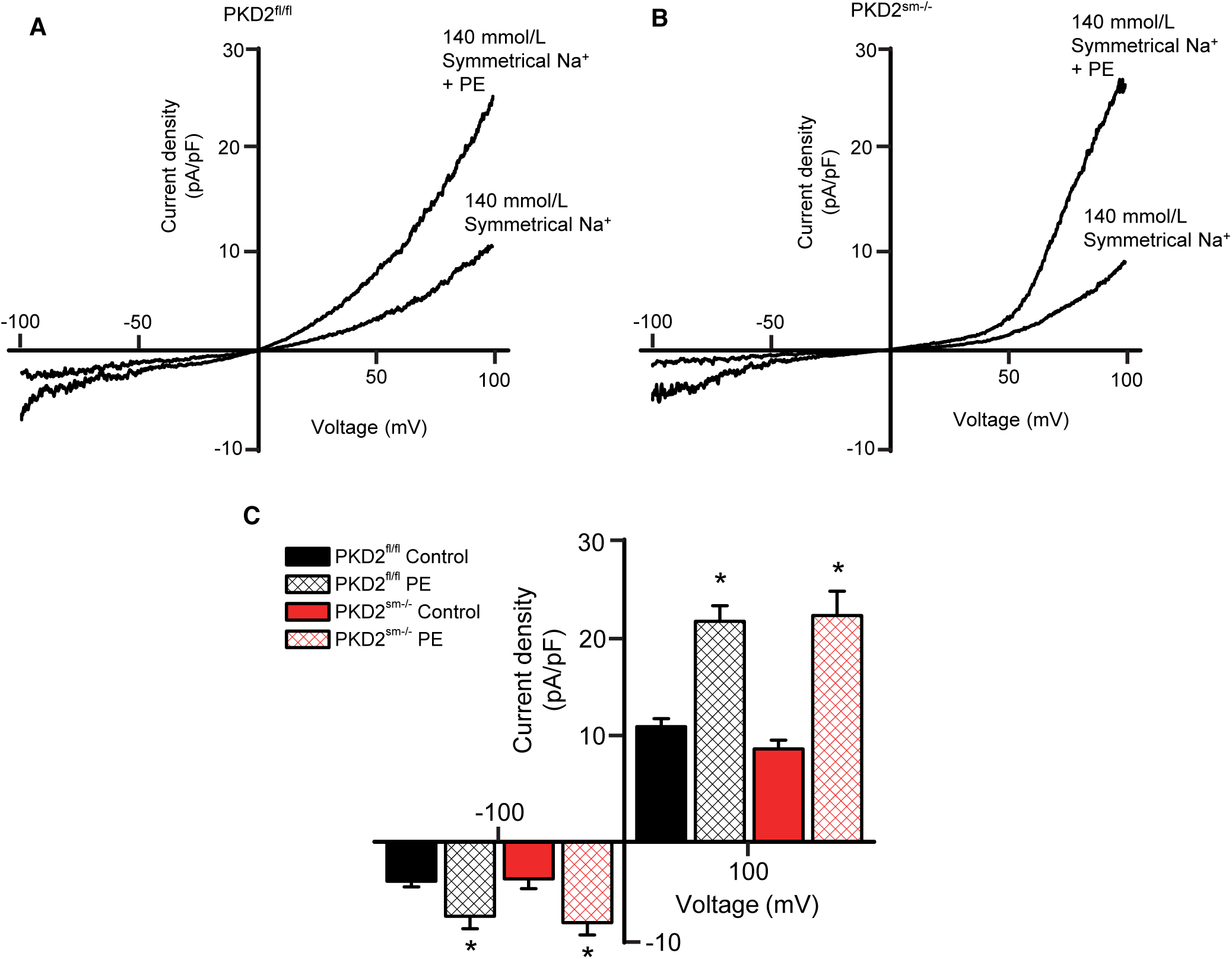
PKD2 knockout does not alter phenylephrine (PE)-activated Na^+^ currents in isolated myocytes from hindlimb artery. A and B: Representative I-V relationships performed in the same isolated hindlimb artery myocytes of PKD2^fl/fl^ or PKD2^sm−/−^ mice in control and PE (10 μmol/L). **C**: Mean data for current density at −100 and +100 mV (PKD2^fl/fl^, n=6 and PKD2^sm−/−^, n=6). * indicates p<0.05 vs control.

**Supplemental Figure 7:**
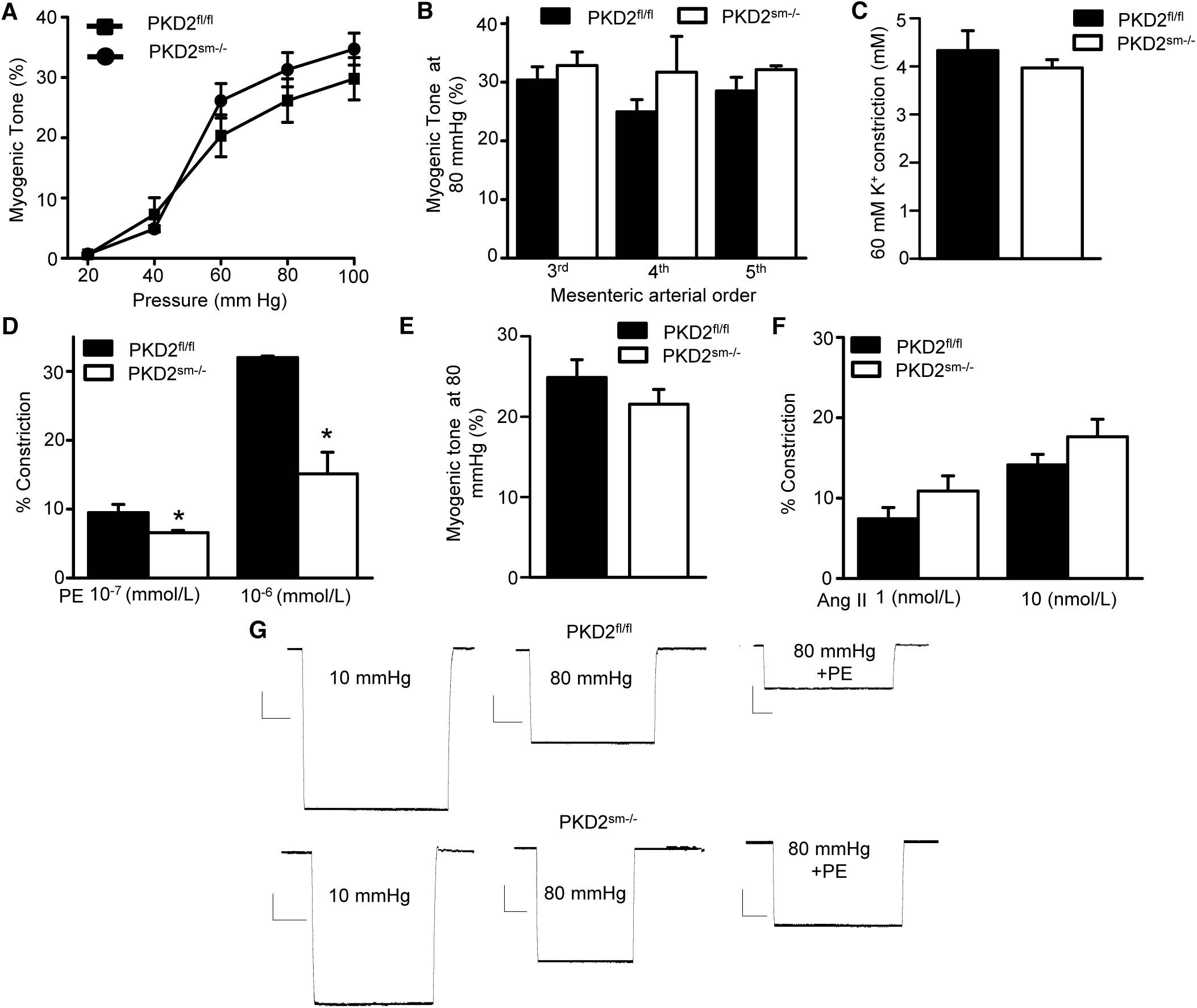
Myocyte PKD2 knockout attenuates phenylephrine-induced membrane depolarization and vasoconstriction, but does not alter pressure or angiotensin II-induced vasoconstriction in hindlimb arteries. **A:** Mean vasoconstriction over a range of pressures in resistance-size mesenteric arteries from PKD2^fl/fl^ (n=7) and PKD2^sm−/−^ (n=9) mice. **B.** Mean myogenic tone at 80 mmHg illustrating that myogenic tone is similar in third-, fourth- and fifth-order mesenteric arteries and unaltered by PKD2 knockout (PKD2^fl/fl^: 3^rd^ n=4; 4^th^ n=5; 5^th^ n=4 and PKD2^sm−/−^: 3^rd^ n=7; 4^th^ n=4; 5^th^ n=4). **C:** Mean data for 60 mmol/L K^+^-induced constriction in first-and second order mesenteric artery rings (PKD2^fl/fl^ n=4; PKD2^sm−/−^ n=5). **D**: Mean data for phenylephrine (PE)-induced vasoconstriction in endothelium-denuded 4^th^ order mesenteric arteries (PKD2^fl/fl^, n=3 and PKD2^sm−/−^, n=3). * indicates p<0.05 vs PKD2^fl/fl^. **E**: Mean myogenic tone at 80 mmHg in endothelium-denuded 4^th^ order mesenteric arteries (PKD2^fl/fl^, n=3 and PKD2^sm−/−^, n=3). **F**: Mean data for angiotensin II-induced vasoconstriction in 4–5^th^ order mesenteric arteries (PKD2^fl/fl^, n=12 and PKD2^sm−/−^, n=10). **G**: Representative traces of microelectrode impalements illustrating that phenylephrine (PE, 1 μmol/L)-induced depolarization is attenuated in arteries from PKD2^sm−/−^ mice. Scale bars: Y= 10 mV, X=20 s.

**Supplemental Figure 8:**
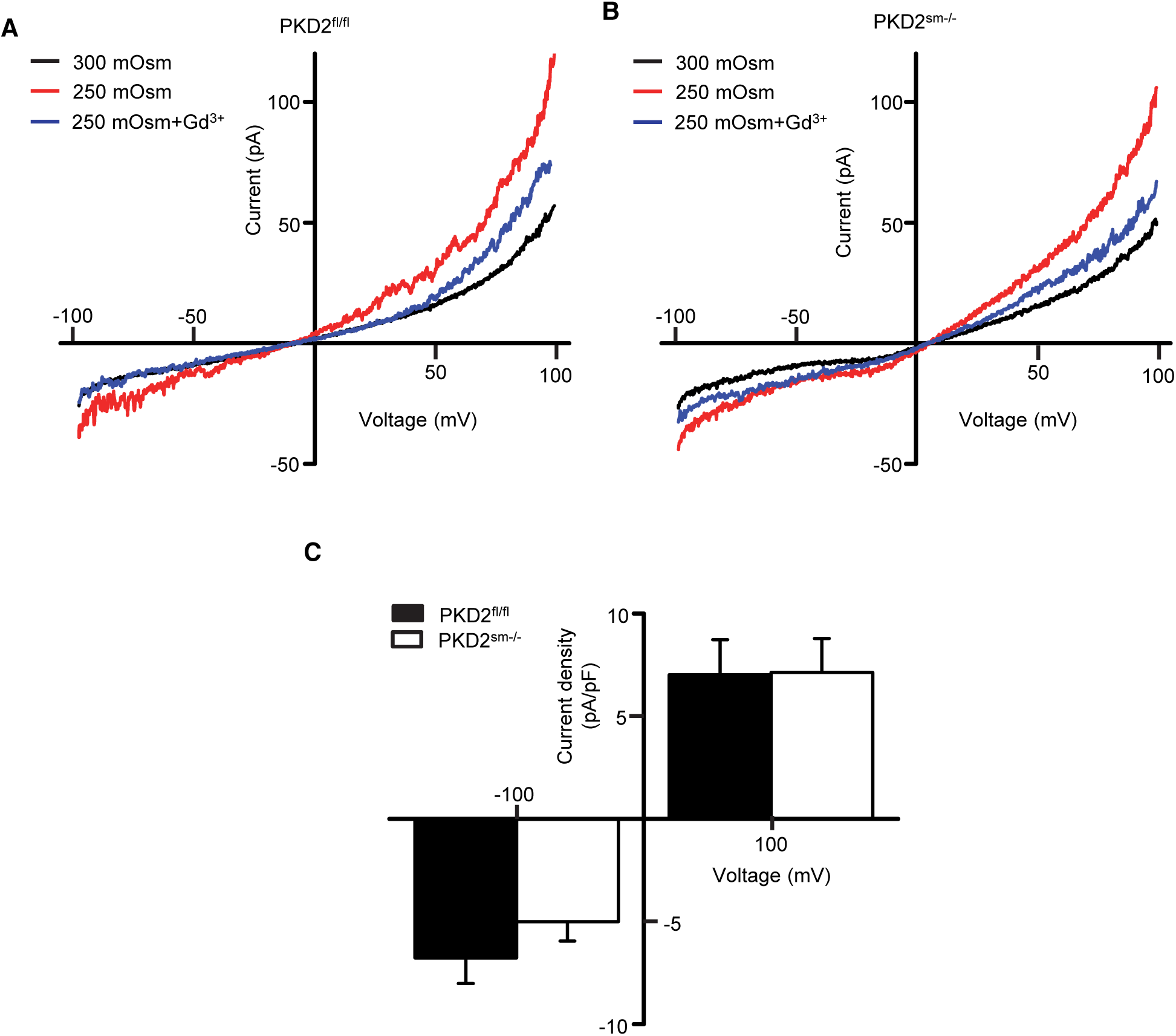
PKD2 knockout does not alter swelling-activated Na^+^ currents in isolated mesenteric artery myocytes. **A and B:** Representative I-V relationships from the same isolated mesenteric artery myocytes of PKD2^fl/fl^ or PKD2^sm−/−^ mice in isosmosmotic (300 mOsm), hyposmotic (250 mOsm) and hyposmotic (250 mOsm) + Gd^3^+ (100 μmol/L). **C**: Mean data for hyposmotic-activated-Gd^3+^ (100 μmol/L)-sensitive cationic current density at −100 and +100 mV (PKD2^fl/fl^, n=6 and PKD2^sm−/−^, n=6).

